# Creased ciliary flocks shape unfolding dynamics via information bottlenecks in an aneural animal

**DOI:** 10.1101/2024.09.30.615941

**Authors:** Charlotte M. Brannon, Manu Prakash

## Abstract

Multicellular organisms utilize thin sheet folding to achieve functional three-dimensional forms. During embryonic development, stereotypical epithelial folds emerge from active cellular and molecular processes including cell shape change and differential cell growth. Active thin sheet folding promises to be a powerful design technique in the fields of active solids, soft robotics, and synthetic biology. However, the general principles of active thin sheet folding remain poorly understood. Here we discover a non-canonical cilia-driven thin sheet folding behavior exhibited by basal animal *Trichoplax adhaerens*. Through volumetric imaging, we found that, despite having no nervous system, *T. adhaerens* has the remarkable ability to resolve complex body folding states in a non-stereotypical fashion using a carpet of collectively flocking cilia. Cilia-resolved imaging revealed that folds create crease defects in the animal’s ciliary carpet, which act as information bottlenecks to break collective behavior of cilia. In turn, these bottlenecks enable the disjointed locomotion required for fold removal. These findings point to a two-way coupling mechanism, wherein ciliary activity shapes the animal’s folding state and vice versa. Our work demonstrates the broad configuration space of non-stereotypical active folding and highlights the power of distributed activity to drive folding and unfolding of a thin multicellular sheet. We anticipate our study to be a starting point for the establishment of a new class of distributed active origami wherein fold lines themselves are dynamic and motile, with implications in engineering of self-folding materials. Additionally, our work reveals a new facet of the Placozoan behavioral repertoire, which extends our understanding of mechanical intelligence in the absence of a nervous system.

## 1. Introduction

Epithelial folding is ubiquitous in animal embryogenesis [1–3] and has been studied in a range of model systems, such as *Drosophila* ventral furrow formation [4] and mammalian intestinal crypt development [5]. Key cellular and molecular processes have been identified which drive developmental epithelial folding, including cellular shape change [6], myosin-dependent apical constriction [7, 8], differential cell growth [9, 10], and more recently, interaction with boundary conditions such as extracellular matrix (ECM) [11] and spatial confinement [12]. These mechanisms, in combination with morphogenic regulation, enable the initiation of folds in a stereotypical, spatiotemporally controlled fashion critical for the robust coordination of developmental processes such as neurulation and gastrulation [13]. Deviations from canonical folding in human development can lead to conditions such as polymicrogyria [14] and spina bifida [15].

Outside the context of animal embryogenesis, diverse examples of multicellular sheet folding can be observed across the tree of life, including in multicellular choanoflagellates [16], bacterial biofilms [17], and *Volvox* embryos [18]. Whereas developmental epithelial folding has been classically understood as a process tightly programmed and controlled at the molecular level [13], these non-canonical examples highlight the general nature of sheet folding in multicellular biology, and demonstrate the capacity for diverse sheet folding processes to emerge from distributed mechanics. Many studies have considered the mechanics of stereotypical epithelial folding [19], but the evolution of complex, non-stereotypical folds in living systems and the broader phase space of epithelial sheet folding has rarely been discussed.

The physics of passive thin sheet folding and crumpling has been studied in a variety of contexts, for example in paper and thin polymer films [20–28]. Yet, how to integrate this body of work with the active self-folding phenomena observed in biology is not obvious. The emerging field of active origami has demonstrated the potential of simple, self-folding robotics, often reliant on active hinges with preprescribed locations [29]. However, these examples lack the distributed activity characteristic of biological systems. Though widespread in biology, folding driven by distributed activity presents key design and coordination challenges, namely: 1) the basic design principles for a self-folding sheet with distributed activity are not understood; 2) the configuration space of possible active thin sheet self-folding forms has not been explored; and 3) the possible coordination rules for distributed active agents controlling self-folding remain unknown.

Here we address these gaps through the discovery of a remarkable thin sheet folding behavior driven by distributed active mechanics in aneural animal *Trichoplax adhaerens* (phylum Placozoa). Spatiotemporal mapping of body folds exhibited by *T. adhaerens* reveals how substrate adhesion and ciliary activity encode the emergent, robust, non-stereotypical folding and unfolding of a thin multicellular sheet. Utilizing a synthesis of imaging methods, including whole-animal tracking microscopy, 4D light sheet fluorescence microscopy, and cilia-resolved time lapse microscopy, we demonstrate that *T. adhaerens* exhibits cilia-driven active folding and unfolding behavior on fast timescales (*∼* seconds-minutes) (SI Movie 1). Using a combination of experimental and theoretical approaches, we discover that folding and unfolding behavior emerges from compartmentalization of ciliary flocks. To the best of our knowledge, this cilia-driven unfolding mechanism represents a novel class of active matter in which folds act as information bottlenecks in a ciliary flock.

## 2. Results

Often described as the simplest extant animal in terms of morphology, *T. adhaerens* displays a flat, sheet-like body composed of fewer than ten morphologically described cell types organized into a dorsal epithelium, intermediate fiber cell layer, and ciliated ventral epithelium responsible for the animal’s locomotion [30, 31]. Despite lacking any musculature or nervous system, *T. adhaerens* is capable of surprisingly complex behaviors [31, 32] derived from both chemical signaling and mechanics [31–33]. Previous studies of *T. adhaerens* have primarily focused on the animal’s flat, surface-attached state [30, 34–36]. Although direct observation of *T. adhaerens* in its natural environment has remained a challenge, prior ecological studies have reported the animal living on algal mats and rocks, which can exhibit complex topographies [37–39]. Furthermore, sampling from the water column implies the existence of a pelagic, surface-detached state [40], which has not been characterized. Motivated by these environmental observations, we investigated the animal’s morphology as a function of substrate context.

### 2.1 Robust, non-stereotypical thin sheet folding and unfolding in aneural animal *T. adhaerens*

By varying local substrate context, we found that *T. adhaerens* can configure into highly complex, three-dimensional folding states (Fig. 1a-c, 1e-h, SI Movie 2). Folding is triggered upon substrate detachment (Fig. 1d), when the absence of a rigid substrate leads to sheet bending and self-adhesion [37, 41]. Folds persist in the body during settling and re-attachment until they are gradually removed through a characteristic “unfolding” behavior (Fig. 1d, SI Movie 1). Notably, re-attachment and flattening behavior has been previously observed in *T. adhaerens* [33, 34, 37, 41–43], but never studied. Moreover, how an aneural organism coordinates a complex series of unfolding steps at the scale of the entire animal is not obvious.

**Fig. 1:**
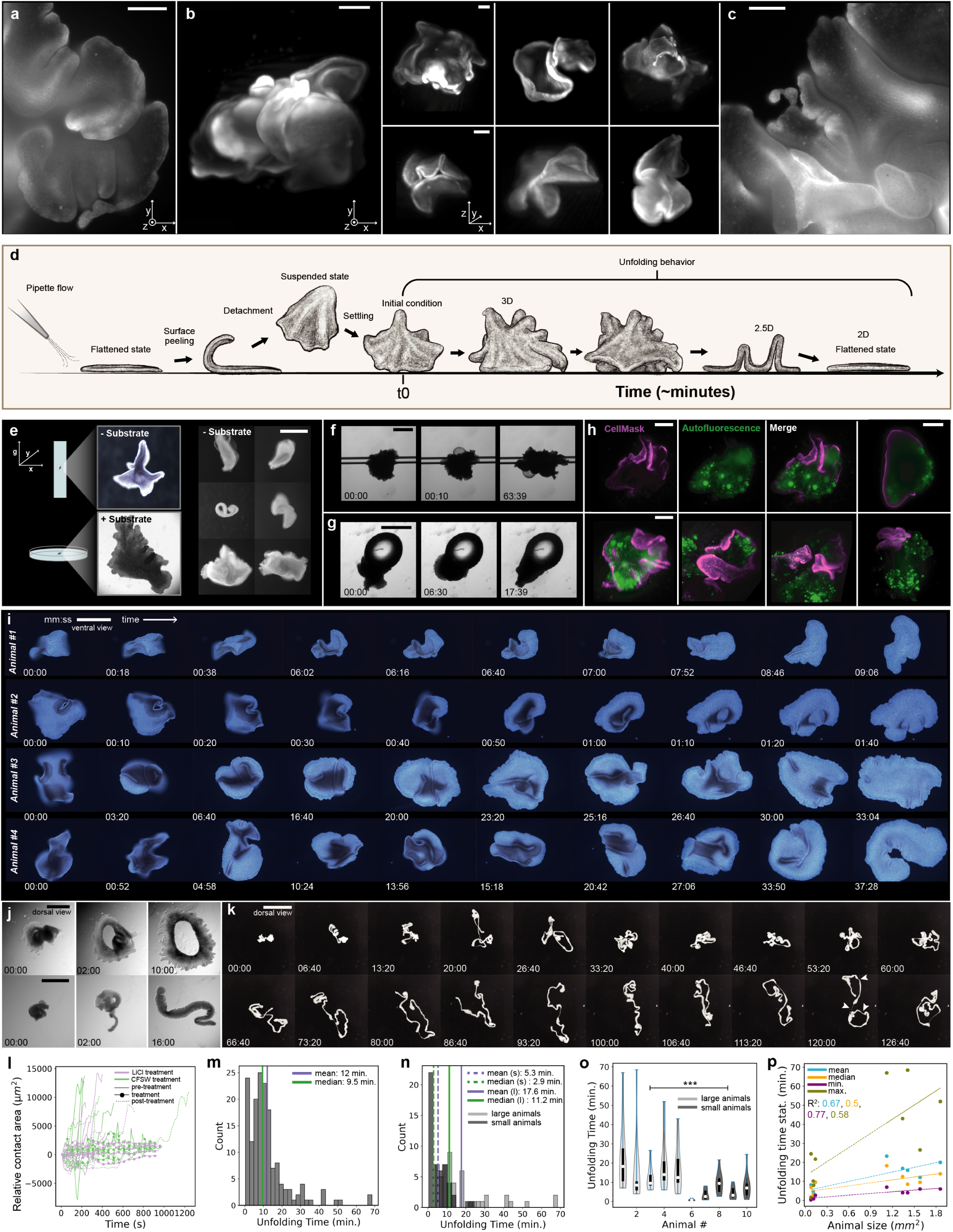
Robust, non-stereotypical thin sheet folding and unfolding in aneural animal *T. adhaerens*. (a-c) Folded states exhibited by *T. adhaerens*, visualized in live animals labeled with membrane stain via (a,c) 2D widefield and (b) 3D light sheet microscopy. Folded states were imaged after post-detachment settling, as shown in panel d. See SI Movie 2. Animals were labeled with CellMask plasma membrane stain. (d) Schematic of the experimental assay used for studying unfolding behavior. We first trigger folding by detaching the animal from a flat, rigid substrate using flow from a pipette. The detached, suspended animal then falls back to the substrate with an initial folding state at *t*_0_, at which point it begins a characteristic “self-unfolding” process to navigate back to a flattened state. (e-h) Impact of substrate context on *T. adhaerens* folding state. (e) In the absence of a rigid substrate, the animal remains in a folded state. Contact with a rigid substrate is necessary for unfolding to occur. See SI Movie 3. We also tested attachment to a range of curved substrates including (f) ultra-thin glass capillaries (170 *µm* diameter), (g) glass microspheres (120 *µm* diameter), and (h) spherical colonial algae, on which the animal naturally feeds. We find that *T. adhaerens* is able to attach and conform to these curved surfaces, maintaining an attached and dynamically evolving folded state. In (h), algae is shown in green (autofluorescence) and the body of *T. adhaerens* is shown in magenta (membrane stain). See SI Movies 4-5. (i) Time lapses of four different animals performing unfolding behavior. Each animal exhibits a different initial condition (far left column), due to random reconfiguration during settling. Each animal passes through a unique, non-stereotypical set of folding intermediates before reaching a flattened state (far right column). (j-k) Animals exhibiting atypical and extreme morphologies are still capable of unfolding, which highlights the general and distributed nature of the animal’s unfolding algorithm. For example, (j) toroidal (“donut shaped”) animals and high-aspect ratio (“stringy”) animals can still undergo complete unfolding. See SI Movie 7. (k) As an extreme case, ultra-long (*∼* 0.5 cm), string-like animals can unfold substantially, though still contain twist in the final state observed (arrows at 120:00 time point). See SI Movie 8. (l) Treatment of animals with calcium-free sea water or 200 mM lithium chloride reversibly inhibits unfolding. Total contact area is used as a readout of unfolding behavior, as the flattened state exhibits the maximum contact area. (m) Histogram of *T. adhaerens* unfolding times for 167 trials across 45 animals demonstrating the robustness of the unfolding algorithm. We measure a mean unfolding time of 12 minutes. (n) Subset of data shown in (m) organized by animal size. Large animals (shown in light grey) consistently take longer to unfold than smaller animals (shown in dark grey. (o) Distribution of unfolding times in (n) sorted by animal identity (labeled 1 to 10). Individual animals exhibit a wide range of unfolding times across repeated trials, and larger animals take significantly longer to unfold than smaller animals (Welch’s *t* (61.07)=5.41, p=1.08e-06). (p) Positive correlation between mean (*R*^2^ = 0.67), median (*R*^2^ = 0.5), minimum (*R*^2^ = 0.77), and maximum (*R*^2^ = 0.58) unfolding times and animal size, further demonstrating the impact of animal size on unfolding time. Scale bars are (a, b - right panels, c, g) 100 *µ*m, (b - left panel) 150 *µ*m, (e) 200 *µ*m, (d) 400 *µ*m, (f, i, j) 500 *µ*m, and (k) 2 mm.

Given that detachment from a flat, rigid substrate triggers folding and re-attachment leads to unfolding, we first investigated the role of substrate geometry in guiding the animal’s folding state. Using vertical tracking microscopy [44], we imaged *T. adhaerens* in the absence of any rigid substrate for prolonged periods (*∼* 30 minutes - 1 hour). Surprisingly, we observed the continued persistence of a folded state, which suggests that attachment to a rigid substrate is necessary for unfolding behavior to occur (Fig. 1e, SI Movie 3). Additionally, we regularly observe folded, detached animals in our cultures, which further supports this intuition (SI Fig. S1). We next tested attachment to curved substrates. Unfolding trials on glass microspheres and thin glass capillaries revealed the animal’s tendency to remain in an attached, dynamically evolving folded state in the presence of high curvature and low substrate area (Fig 1f-g, SI Movie 4). Fortuitously, we discovered a similar behavior occurring naturally in our cultures, where animals were observed attaching to, folding on, and possibly feeding on roughly spherical algae clusters (sp. unknown) (Fig. 1h, SI Movie 5). Previously undescribed, this behavior may represent a novel physiological mode of *T. adhaerens* in the ocean. Moreover, these cases of attachment to irregular substrates exhibited by this ultra-thin animal highlight the high-adhesion, low-bending energy regime [26], which leads to folding or unfolding depending on substrate geometry.

Next, to test the robustness of unfolding behavior and quantify unfolding pathways, we collected 163 unfolding time lapses (on flat substrates) from repeated trials across 45 animals (Fig. 1d, 1m). This dataset reveals unfolding behavior to be highly robust; all trials resulted in the animal reaching a final flattened state, except for one which ended in animal fission at the site of a twist (SI Movie 6). From these data, it is apparent that unfolding behavior is highly non-stereotypical; each animal passes through a series of distinct folding intermediates (Fig. 1i). Interestingly, this is also the case for individual animals throughout repeated unfolding trials (SI Fig. S2, S3). Thus, unlike developmental folding, which is highly stereotypical [45–48], we find that *T. adhaerens* folding and unfolding behavior is far more varied and unpredictable, as if the animal is solving a folding problem *ad libitum*. Furthermore, the non-stereotypical, fast nature of *T. adhaerens* unfolding suggests the likely role of mechanics, rather than genetics, in driving the behavior, which prompts us to wonder: what possible distributed “mechanical algorithms” can enable robust unfolding of a thin multicellular active sheet?

It is well known that *T. adhaerens* exhibits extreme (clonal) variation in body size and shape [49]. To test the impact of animal shape on unfolding, we performed unfolding trials for toroidal animals and high-aspect ratio “stringy” animals [49], which are some of the less common (though naturally occurring) animal morphotypes. We find that toroidal animals can still unfold completely despite their unusual body plan (Fig. 1j, SI Movie 7). Interestingly, high-aspect-ratio animals are also capable of significant unfolding (Fig. 1j-k, SI Movie 8), but after two hours of unfolding, twist defects persist in the body (Fig. 1k, arrows). Remaining twist defects often lead to binary fission of the animal. These findings highlight the general and distributed nature of unfolding behavior, which can run on a wide range of body morphologies.

It naturally follows to consider the impact of animal size on unfolding. Using a mixture of small and large animals (varying by roughly one order of magnitude in area), we performed ten repeated unfolding trials across ten animals (100 total measurements) (Fig. 1n). We find that, on average, large animals are roughly three times slower to unfold compared to small animals, likely due to their ability to configure into more complex folding states (Fig. 1n-p) [50]. Additionally, organizing unfolding times based on animal identity reveals a wide range of unfolding times in repeated trials, particularly for larger animals, which correspond to variations in unfolding paths taken (Fig. 1o, SI Fig. S2, S4). Aside from these findings, unfolding behavior is robust to animal size.

Finally, previous work has demonstrated the role of the animal’s ventral ciliary carpet, which exhibits characteristic collective flocking dynamics, in driving locomotion [31, 51]. We tested whether normal ciliary activity is necessary for unfolding behavior by performing unfolding trials in calcium-free seawater and lithium chloride, both of which have been previously shown to perturb normal ciliary beating in *T. adhaerens* [43] (Fig. 1l, SI Fig. S5, see Methods). As predicted, we find that both treatments reversibly inhibit unfolding, which demonstrates the critical driving role of ciliary activity in unfolding behavior.

### 2.2 3D structural characterization of folding states

Having shown that unfolding behavior is an active, non-stereotypical folding process dependent upon local substrate context, we next aimed to study this behavior in 3D. Previous studies in the fields of thin sheet folding and crumpled paper physics have utilized volumetric imaging techniques to establish the relationship between three-dimensional structure and folding/unfolding mechanism [21, 52–54]. Thus, to investigate the unfolding mechanism at play in *T. adhaerens*, we performed 4D light sheet and 3D confocal scans of both fixed and live fluorescently labeled animals undergoing unfolding behavior (Fig. 2, SI Movie 9, see Methods).

**Fig. 2:**
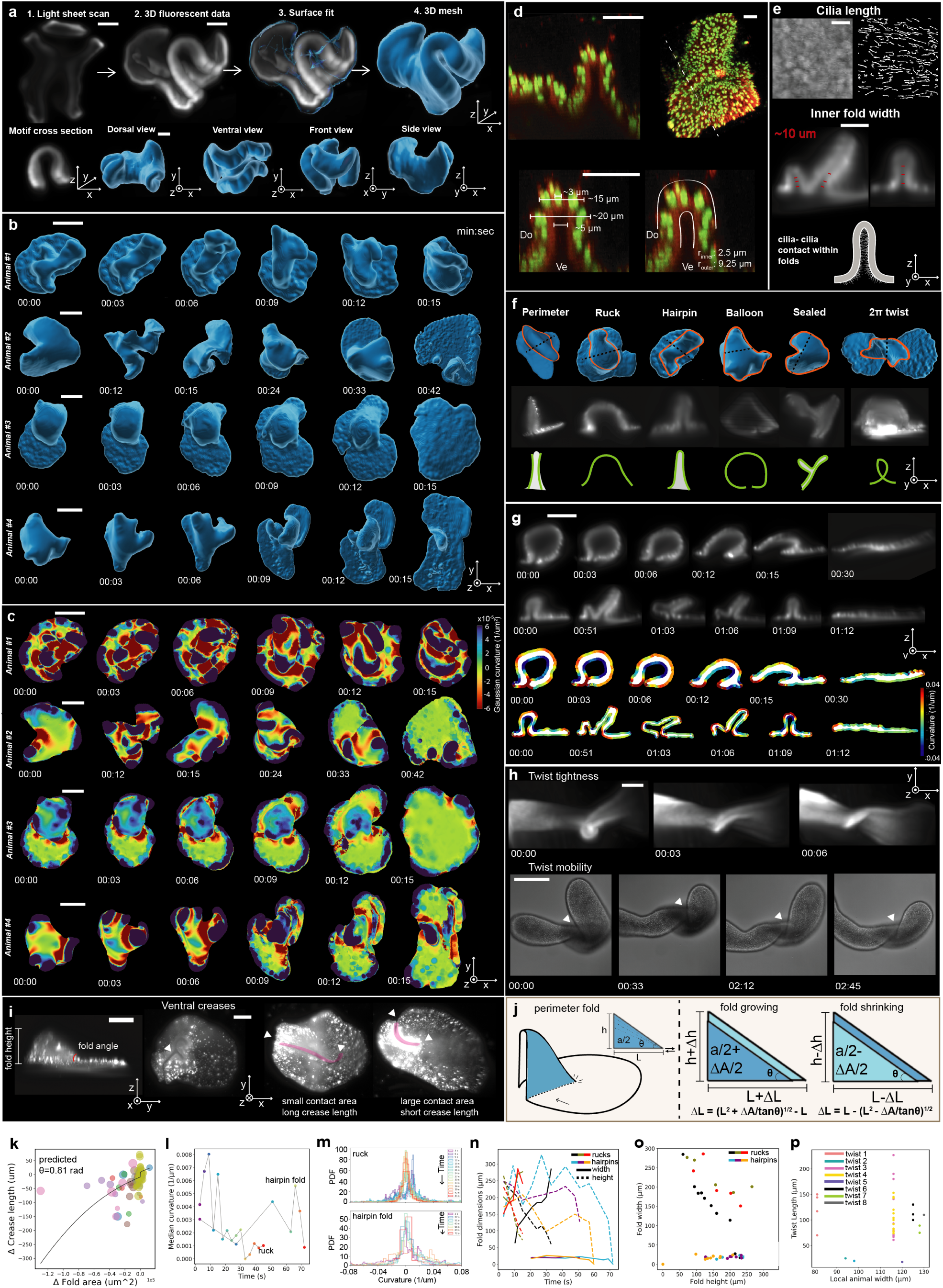
3D structural characterization of folding states. (a) 3D light sheet microscopy of folded *T. adhaerens*. Animals were stained with either CellMask membrane stain or LysoTracker to enable visualization of the entire body. See SI Movies 9-10. (b) 4D light sheet imaging of the unfolding process in four different animals. Surface meshes shown. The variety of unfolding trajectories and fold structures are apparent. (c) Local Gaussian curvature of meshed 3D states shown in (b). Curvature evolution is non-monotonic, which demonstrates the importance of activity in shaping the animal’s folding state. (d) Cellular perspective within folds. Confocal scans of fixed folded animals labeled with Hoechst nucleic acid stain (green) and CellMask plasma membrane stain (red) reveal *∼* 3 cells spanning the cusp of a high-curvature fold. (e) Extraction of inner fold width and ciliary length (both *∼* 10 *µ*m), which implies cilia-cilia contact within folds. (f) Commonly observed folding motifs are perimeter folds, rucks, hairpin folds, balloon folds, sealed folds, and 2*π* twists. (g) Cross sectional time lapse imaging of individual folds and their local curvature maps. (h) 2*π* twists exhibit striking mobility and variation in tightness. Unlike folds, twists behave as point defects which must be removed from the edge of the body. See SI Movie 11. (i) Side view of a perimeter fold enables visualization of a fold height and angle (i.e., the angle the fold base makes with the substrate). Ventral views of perimeter folds reveal crease lines, which vary in length as material is added to or removed from the fold. (j) From a simple geometric representation of a perimeter fold, we estimate how crease length is expected to scale with fold area as folds grow and shrink in height. (k) We extracted fold area and crease length values from nine perimeter folds across different datasets, and fit these data to the equations in panel j. The resulting fit predicts a fold angle of *∼* 45°. See SI for further discussion. Circle size represents crease length. Color represents fold identity. (l) Median curvature evolution for the ruck and fold shown in (g), which further demonstrates that curvature evolution is non-monotonic and varies depending on fold structure. Hairpin folds with self-contact exhibit relatively steady median curvature until the fold is removed. On the other hand, rucks undergo gradual curvature reduction as the ruck is removed. (m) Curvature evolution distributions for the ruck and fold shown in (g). (n) Fold width and height time evolution for the ruck and fold shown in (g) and additional rucks and folds shown in SI Fig. S10. (o) Fold width and height scaling. Hairpin folds (blue, orange, purple) exhibit constant fold width for variable heights, while ruck width scales with height (black, green, red). (p) Quantification of twist tightness. A single twist within a given animal can exhibit highly variable widths, as thin as 10 *µ*m, which is on the order of a single cell diameter. See SI Movie 11. Scale bars are: (a, c bottom row, h top row) 100 *µ*m, (b) 20 *µ*m, (d, e, g) 200 *µ*m, (c top row) 10 *µ*m, and (h bottom row) 500 *µ*m. Axes show image orientation; we define the z axis as the normal to the substrate.

From these data, it is immediately apparent that folds are high curvature, hierarchical structures, which display highly variable geometries (Fig. 2a-c, SI Movie 9). Folds are distributed throughout the animal’s body, and do not leave a structural memory after being resolved (unlike crumpled paper) (Fig. 2b). Furthermore, folding states are highly dynamic and exhibit dramatic changes on the order of seconds (Fig. 2b). We fit smooth surfaces to these folded geometries and extracted local Gaussian curvature across the body (Fig. 2c, SI Movie 10, see Methods), which revealed the animal’s ability to increase body curvature in the process of unfolding before eventually reaching a flat state. The non-monotonicity of curvature evolution highlights the inherent frustration in the associated energy landscape for unfolding (Fig. 2c). To determine the cellular length scale relative to fold curvature, we performed confocal scans of fixed folded animals, which strikingly revealed, in the most extreme case, three cells spanning the fold cusp (Fig. 2d). Additionally, we extracted an inner fold width of 5-10 *µm*, which implies some degree of cilia-cilia contact within high-curvature folds (Fig. 2e) [31].

Previous studies of animal morphogenesis have revealed distinct epithelial folding motifs such as furrows, ridges, branches, and loops [55]. Although *T. adhaerens* exhibits a wide variety of non-stereotypical folding states (Fig. 2b), we outline common folding motifs, shown in Figure 2f, as follows: 1) perimeter folds, 2) rucks [24], 3) hairpin folds, 4) balloon folds, 5) sealed folds, and 6) 2*π* twists. Interestingly, all motifs (except for 2*π* twists) are biased toward negative ventral (positive dorsal) curvature, which is likely indicative of the role of adhesion between the ventral cilia and substrate in driving folding and unfolding. Tracking the time evolution of hairpin folds and rucks (Fig. 2g) reveals variable scaling between height and width of these structures (Fig. 2n-o). We find that ruck height scales with ruck width, while hairpin folds change height without changing width, likely due to adhesive self-contact within the fold (Fig. 2n-o). Additionally, extracting local cross-sectional curvature allows us to quantify motif-specific curvature evolution (Fig. 2g, 2l-m). We find that rucks undergo gradual curvature removal, while hairpin folds maintain constant curvature until the moment they are removed (Fig. 2g, l-m).

We next focus on the scaling relationship between fold area and length. Strikingly, hairpin and perimeter folds display distinct crease lines on the ventral surface, which are observed to change in length over time as more material is absorbed into (or removed from) the fold (Fig. 2i, Fig. 3). From the rough geometry of a growing perimeter fold, we show that the change in length should scale approximately as 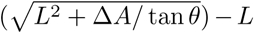 and for a shrinking fold, 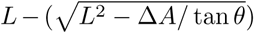, where *θ* is the maximum angle (based on sheet elasticity) the fold base can make with the substrate (Fig. 2i-j, see Methods and SI for details). Interestingly, this behavior is analogous to contact line pinning dynamics in liquid droplets on surfaces, which only begin to expand in contact area upon reaching a critical angle set by the liquid and surface properties [56] (see SI for further discussion). Fitting the data in Figure 2k to the above equations yields a predicted *θ ≈* 45*^◦^*, which is similar to that observed in our 3D data (Fig 2i,2k).

**Fig. 3:**
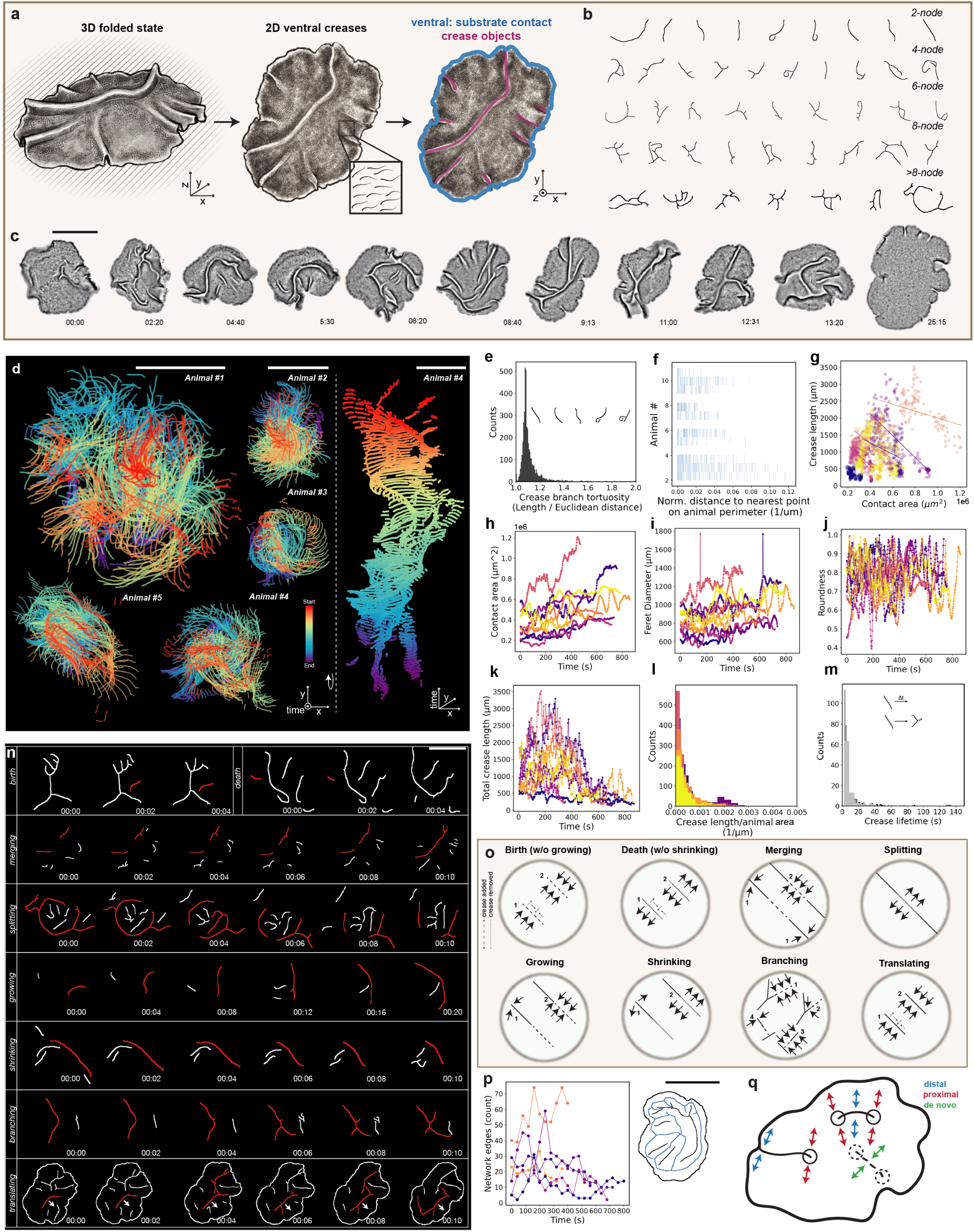
Unfolding dynamics through the lens of ventral crease defects. (a) Imaging of the animal’s ciliated ventral surface during unfolding reveals 2D crease lines corresponding to 3D folds. We consider the time evolution of creases over the course of unfolding. Crease and body contours were extracted from 10 unfolding time lapses across 10 animals. See SI Movie 12. (b) Example crease geometries sorted by node number (i.e. the number of network junctions), ranging from simple linear 2-node creases to more complex 4-, 6-, 8-, and *>*8-node creases. (c) Example unfolding time lapse in which ventral creases are visible. Imaging in low-concentration fluoroscein boosts crease signal, as the dye fills the negative space created by folds. (d) Crease positions over time for five different unfolding animals. Time indicated by color. (e) Tortuosity was calculated for all extracted crease branches, which reveals the prevalence of curved creases. (f) Normalized distance of crease centroids to the nearest point on the animal’s border. Across animals and over the course of unfolding, there is a broad distribution of distances, which reflects the lack of preprescription of crease positions. (g) Negative correlation between crease length and animal contact area. As material is absorbed into folds, crease length increases while animal contact area decreases. Color indicates animal identity. (h) Contact area, (i) Feret diameter, (j) roundness, and (k) total crease length vs. time for ten different unfolding animals. Fluctuations in animal contact area and shape are observed. Both contact area and total crease length evolve non-monotonically. In many animals, total crease length increases significantly before decreasing. Color indicates animal identity. (l) Histogram of crease lengths normalized by animal size for ten unfolding animals. Most creases are short, rather than spanning the entire body, which reflects the inhomogeneous nature of ciliary compression and stretch across the body. (m) Histogram of crease lifetimes observed across ten unfolding animals. Creases are short-lived, typically lasting 5-10 seconds until disappearing or merging with another crease. (n) Unit operations of creases. Creases were observed to undergo birth, death, merging, splitting, growing, shrinking, branching, and translation events. These operations remodel the contact domains of the ventral surface with the substrate. See SI Movie 13. (o) Schematic showing the local tissue motions necessary to perform crease operations shown in (n). (p) Ventral contact domain network analysis highlights the topology of the ventral surface in contact with the substrate (the path outlined by crease patterns). Crease operations shown in (n) lead to remodeling of the ventral surface domains. Color indicates animal identity. (q) Schematic summarizing how different tissue motions can lead to crease remodeling. Both distal (blue) and proximal (red) tissue motions can contribute to crease unit operations such as lengthening and shortening (as discussed in section 2.2). *De novo* motions may also lead to the birth of new creases. Scale bars are 500 *µ*m.

Finally, we consider the 2*π* twist motif. We clearly observe variability in twist tightness within a single animal over the course of unfolding (Fig. 2h, 2p, SI Movie 11). The tightest twist observed in our data is *∼* 10 *µm* wide, which is on the order of a single dorsal epithelial cell diameter. Additionally, time lapse data revealed extensive twist mobility, similar to that of topological solitons or “twistons” as observed in graphene nanoribbons [57]. It is possible that twist geometry, which exhibits liftoff from the substrate, may contribute to its mobility. Both 2*π* and *−*2*π* twists can form in the same animal, enabling twist annihilation and pileup, depending on twist chirality (SI Fig. S6). Twist has been explored in a variety of contexts [58–62]. In later sections, we further explore the impact of twist on cellular physiology and implications for unfolding behavior.

### 2.3 Unfolding dynamics through the lens of ventral crease defects

Having characterized the folding states of *T. adhaerens* in 3D, we next turned our attention to the animal’s ventral surface in order to investigate the role of ciliary activity in unfolding behavior from a mechanistic perspective. We collected unfolding time lapses of the animal’s ventral surface in low-concentration fluorescein (see Methods), which enabled the segmentation and analysis of animal contact area contours and 2D crease defects corresponding to the base geometry of 3D folds (Fig. 3a-d, Fig. 2i, SI Movie 12). This low-dimensional approach allows the observation of statistics and rules governing unfolding, and invites us to consider how substrate contact domains of active ciliary patches shape the positioning of folds and vice versa. We first extracted all creases from ten different unfolding time lapses taken from ten different animals (Fig. 3a-d). We find that creases range from simple line defects (i.e. 2-node, Fig. 3b) to complex network-like structures (4-, 6-, 8-, *>*8-node, Fig. 3b). Analysis of crease tortuosity (ratio of path length to displacement) reveals a mean tortuosity *>* 1, which highlights the ability of local ciliary activity to shape crease geometry (Fig. 3e). Additionally, we extracted crease positions relative to the animal perimeter, and observe a broad distribution of positions, which shows the lack of any stereotypical prescription of creases (Fig. 3f).

To shed light on the mechanisms of crease removal, we next focused on crease and contact area dynamics. Growth of a crease, by definition, must “consume” animal area, leading to both height and length increases (see fold length and area scaling discussion in section 2.2), and reduction in total contact area (Fig. 3g, 2i). Contact area is therefore a useful representation of the animal’s folding state. We discover that animal-substrate contact area increases non-monotonically over the course of unfolding (Fig. 3h); cilia-substrate contact domains grow and shrink before eventually flattening, a likely signature of active ciliary dynamics (SI Fig. S8). Furthermore, unfolding is associated with dramatic changes in contact contour shape, as captured via Feret diameter and roundness metrics (Fig. 3i-j). This finding could have interesting implications for ciliary activity during unfolding, since it has been previously suggested that animal shape and size may impact collective ciliary dynamics [51, 63]. Surprisingly, we observe that for many animals, total crease length *increases* before decreasing with unfolding (Fig. 3k), which could imply the possible utility of crease defects in promoting eventual unfolding. Finally, we found that many creases are short relative to the body size (Fig. 3l). Additionally, creases are short lived, with an average lifetime of *∼* 5-10 seconds before being resolved or merging with another crease (see Methods, Fig. 3m). Importantly, the presence of short creases reflects the spatial heterogeneity of compression across the body arising from distributed ciliary activity.

We next tracked the evolution of individual creases in order to determine how the local ventral landscape is remodeled over the course of unfolding. Strikingly, we find that creases are highly dynamic, changing and evolving on timescales of seconds (Fig. 3k, 3n). We observe cases of crease birth, death, merging, splitting, growing, shrinking, branching, and translating (average crease translation speed of *∼* 3 *µm/s* in the animal frame of reference, as shown in SI Fig. S7) (Fig. 3n). Each of the observed crease operations can be understood as a corresponding motion of the adjacent tissue (and therefore cilia [31]), as shown in Figure 3o). Interestingly, crease tips appear to exhibit attraction to one another, which leads to merging and “annihilation” of two crease tips (SI Movie 13). In this way, we liken crease tips to topological defects [64]. Furthermore, these unit operations of crease behavior have interesting consequences for ciliary activity on the ventral surface. As previously mentioned, it has been shown that *T. adhaerens* ventral cilia exhibit collective flocking dynamics mediated by tissue elasticity [51]. However, folds inherently change the local elastic landscape by creating a crease “defect”. Thus, as creases move and change shape, they are simultaneously changing the elastic connectivity of ventral cilia (Fig. 3p). We explore the consequences of these dynamics in section 2.5.

### 2.4 1D scaling analysis and toy model of active folding and unfolding

Having considered the behavioral, structural, and dynamical aspects of unfolding, we next take an energetic approach to understand the configuration space of active sheet unfolding in a 1D model system (Fig. 4a).

**Fig. 4:**
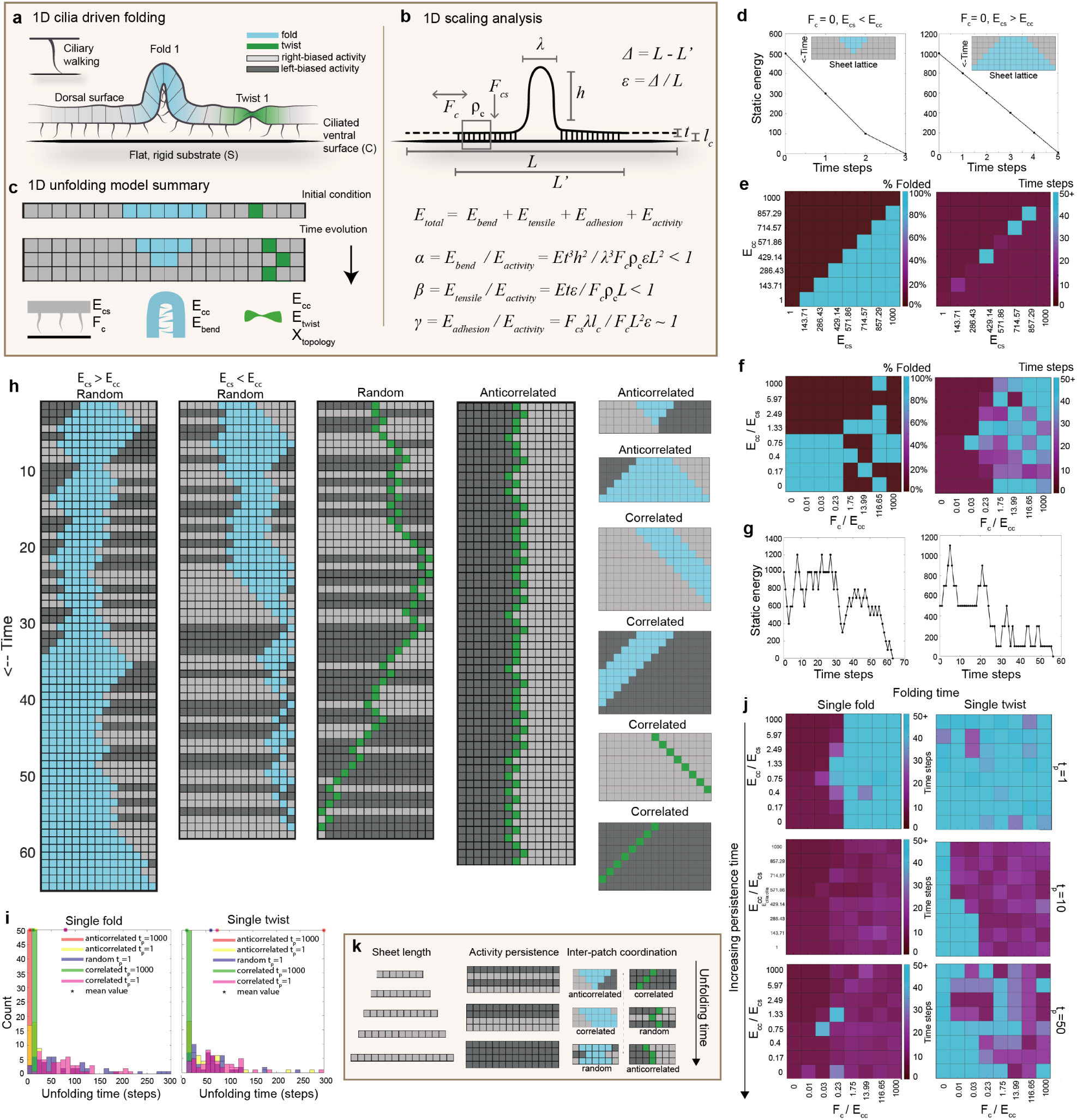
1D scaling analysis and toy model of active folding and unfolding. (a) We consider the energetics of active folding and unfolding in a 1D context. (b) Scaling analysis of active unfolding reveals unfolding to be driven by activity, rather than sheet elasticity. See SI for further details. (c) We develop a 1D toy model to capture the time evolution of hairpin folds, twists, and active flat domains in a 1D lattice. Self-adhesion energy *E_cc_* is associated with folds and twists, while substrate adhesion energy *E_cs_* is associated with flat regions. Activity of flat regions biases the motion of fold and twist boundaries. This scheme enables the consideration of how different activity algorithms impact folding behavior in a simple system. See SI for model details. (d) Sheet evolution with zero activity and either *E_cs_ < E_cc_* (left) or *E_cs_ > E_cc_* (right). These passive regimes lead to unfolding and folding, respectively. (e-f) Folding state and unfolding time phase spaces for sheet evolution with (e) zero activity and (f) nonzero random activity. In (f), when activity is stronger than adhesion energy, activity orientation determine the final folding state at the cost of longer unfolding times. Activity of a patch is oriented to the right or left for a persistence time *t_p_* before randomly reorienting. (g) Example energy vs. time plots for two random fold cases shown in (h). Activity biases the system to reach high-energy states before eventual folding or unfolding. (h) Outcome of naive patch coordination schemes for sheets containing a single twist (green) or single fold (cyan). Random, anti-correlated, and correlated activity lead to variable unfolding outcomes. For example, anti-correlated activity leads to fast fold removal but frustration in the case of twist. (i) Unfolding time distributions for algorithms shown in (h) for folds (left) and twists (right). For folds, anti-correlated activity with long persistence time leads to fastest unfolding times followed by correlated, activity with long persistence time, anti-correlated activity with short persistence time, correlated activity with short persistence time, and random activity. For twists, correlated activity with long persistence time leads to fastest unfolding time followed by random activity, correlated activity with short persistence time, anti-correlated activity with short persistence time, and anti-correlated activity with long persistence time. (j) Impact of persistence time on unfolding time. Low persistence time (*t_p_* = 1 time step, top row) results in long unfolding times; intermediate persistence time (*t_p_* = 10 time steps, middle row) results in reduced unfolding times; and high persistence time (*t_p_* = 50 time steps, bottom row) results in long unfolding times due to the maintenance of frustrated states. (k) Summary schematic. Animal size, cilia persistence time, and patch coordination all impact unfolding time in this 1D system.

The physics of thin sheet folding and deformation has been studied in a variety of contexts [20–27, 65–67], including substrate delamination [24, 67], elastic wrinkling [25, 66], paper crumpling [68], and paper unfolding [23]. These are passive processes in which material properties such as the Young’s modulus, bending modulus, and self- or substrate-adhesion govern the folding state in response to external force (SI Fig. S8). Uniquely in the case of living self folding sheets such as *T. adhaerens*, we must also account for the dynamics of distributed of activity which injects energy into the system at the micron length scale.

To understand where *T. adhaerens* lives in the phase space of active folding, we consider a 1D folded sheet with length *L*, thickness *t*, fold width *λ* and fold height *h* (Fig. 4b) [24, 67]. The strain in this system *ɛ* is given by the length of excess material that has been absorbed into the fold, Δ = *L − L^′^* (Fig. 4b). We define the relevant energy scales in the problem as: 1) *E_bend_* (out-of-plane bending energy), 2) *E_tensile_* (in-plane elastic energy), 3) *E_cilia:substrate_* (adhesion energy between the cilia layer and the substrate, abbreviated as *E_cs_*), and 4) *E_activity_* (energy injected by ciliary walking). Leveraging past measurements of the bending modulus *B ∼* 2 *×* 10*^−^*^13^ *N · m* and Young’s modulus *Et ∼* 1.5 *×* 10*^−^*^2^ *N/m* in thin epithelial sheets [27], we evaluate the Föppel-von Kármán number *γ* = *Y L*^2^*/B ∼* 7 *×* 10^4^, which is much larger than 1 (see Methods and SI for details). Thus, out-of-plane bending is the preferred mechanism for releasing in-plane stress, which leads to the ready formation of folds in the system (Fig. 3). Additionally, we can write the total energy in the system as: *E_total_* = *E_bend_* +*E_tensile_* +*E_cs_* +*E_activity_*. *E_cs_* = *F_cs_ρ_c_λl_c_*, where *F_cs_* is the adhesion energy per cilium, *ρ_c_* is the ciliary density, and *l_c_* is the ciliary length; *E_bend_* = *Et*^3^*h*^2^*/λ*^3^; *E_tensile_* = *Et*^3^*ɛ*^2^*L*, where *ɛ* = Δ*/L*; and *E_activity_* = *F_c_ρ_c_ɛL*^2^, where *F_c_* is the force exerted by a walking cilium. Using the above values for *E_bend_* and *E_tensile_* [27]; previous measurements of cilia-substrate adhesion *F_cs_ ∼* 2 *nN/*cilium [69]; previous estimates of active ciliary force *F_c_ ∼* 10 *pN* [70, 71]; and measurements of ciliary density *ρ_c_ ∼* 0.4 cilia*/µm* (see Methods), we evaluate *α* = *E_bend_/E_activity_ ∼* 0.158, *β* = *E_tensile_/E_activity_ ∼* 0.06, and *γ* = *E_cs_/E_activity_ ∼* 2.17 (see SI for details). Thus, active ciliary forces dominate over elastic forces, which permits the emergence of creases as a function of local ciliary activity (Fig. 3). Notably, ciliary adhesive and active forces are comparable in magnitude, but because active forces scale with *L* (which can be large or small) and adhesive forces scale with *λ* (which is expected to be invariantly small and invariant, as seen in Fig. 2-3), unfolding behavior is largely driven by active rather than passive properties of the sheet (Fig. 4b).

While scaling-based calculations capture the possibility of folding in active elastic sheets, they do not elucidate a kinetic path for folding/unfolding dynamics. Thus, we develop two reduced order discrete models that represent cells as lattice sites, each with an associated activity: 1) a 1D toy model to explore possible cilia coordination schemes during unfolding (Fig. 4) and 2) a 2D model to investigate how unfolding behavior emerges from ciliary flocking dynamics (section 2.6, Fig. 6). Sequentially adding complexity in this manner allows us to search for mechanical algorithms and associated robustness in 1D and 2D active sheet folding systems.

We consider a simple 1D toy model of a string of cells with in-plane stiffness *E_tensile_*, bending modulus *E_bend_*, surface adhesion *E_cs_*, ciliary activity *F_c_* (see SI for details), and self-adhesion *E_cc_*, which represents the adhesion energy between a pair of cilia in contact within a hairpin fold (see Methods, SI, Fig. 4c). Behavior of this string is set by a competition between *E_cs_*, *E_cc_*, and *F_c_* (Fig. 4c). For simplicity, we consider how such a discrete sheet behaves with two simple folding motifs, hairpin folds and 2*π* twists (Fig. 2f). In the passive case (*F_c_* = 0), folding or unfolding occurs based on the ratio *E_cs_*/*E_cc_* (Fig. 4d-e). When (*F_c_ >* 0), active forces (arising from flat regions of the lattice) can directly influence the folding state (Fig. 4f-g).

Using this 1D toy model, we tested different coordination schemes for distinct regions of flat cilia and their impact on final folding state and unfolding time for single-fold and single-twist initial conditions (Fig. 4h-i). In the random case, distinct flat regions cannot communicate with one another, and are randomly reoriented every *t_p_* time steps (where *t_p_* is defined as the persistence time). This scheme is capable of removing both folds and twists, though inefficiently (Fig. 4h-i). In the anti-correlated case, flat regions “know” the directionality of their neighboring region and take the opposite direction (Fig. 4h). This scheme can successfully remove folds but leads to frustration in the case of twist and vice versa in the case of flipped directionality (i.e., the left patch exhibiting rightward activity and right patch exhibiting leftward activity, not shown). Finally, correlated regions (i.e., all patches demonstrate activity in the same direction) can push both folds and twists out of the lattice, though a velocity differential between the two regions is required for this to occur (see SI for further discussion). By running 50 simulations with the same initial folding condition (single fold or single twist) for each coordination scheme, we collected unfolding time statistics revealing marked performance differences for different algorithms when comparing single-fold and single-twist systems (Fig. 4i). Importantly, a fold may be removed in its place while a twist must be pushed to the boundary of the sheet due to its topology.

Next we considered the role of sheet size and activity persistence time in 1D random unfolding (as defined above). We find that for a given fold size *h*, larger sheet sizes exhibit longer unfolding times due to an increased likelihood of nucleating new folds based on a greater number of lattice sites (SI Fig. S9). For a given sheet *L* and fold size *h*, low persistence time leads to long unfolding times for both single-fold and single-twist lattices (Fig 4j, top row). Increasing persistence time leads to a reduction in unfolding times (Fig 4j, middle row). However, very high persistence time leads to the re-emergence of long unfolding times, as frustrated states are maintained for at least *t_p_* time steps (Fig 4j, bottom row). Thus, there is a fundamental trade-off between fold removal speed and frustration risk. Consequently, expected fold height should influence the choice of an optimal persistence time (e.g. smaller folds require shorter persistence times).

Importantly, real ciliary activity in *T. adhaerens* exhibits far more complex flocking dynamics [51] than can be considered in a 1D lattice. Therefore, we next performed experiments to identify cilia flocking dynamics at play during unfolding behavior (section 2.5, Fig. 5), and designed 2D simulations to capture these dynamics (section 2.6, Fig. 6).

**Fig. 5:**
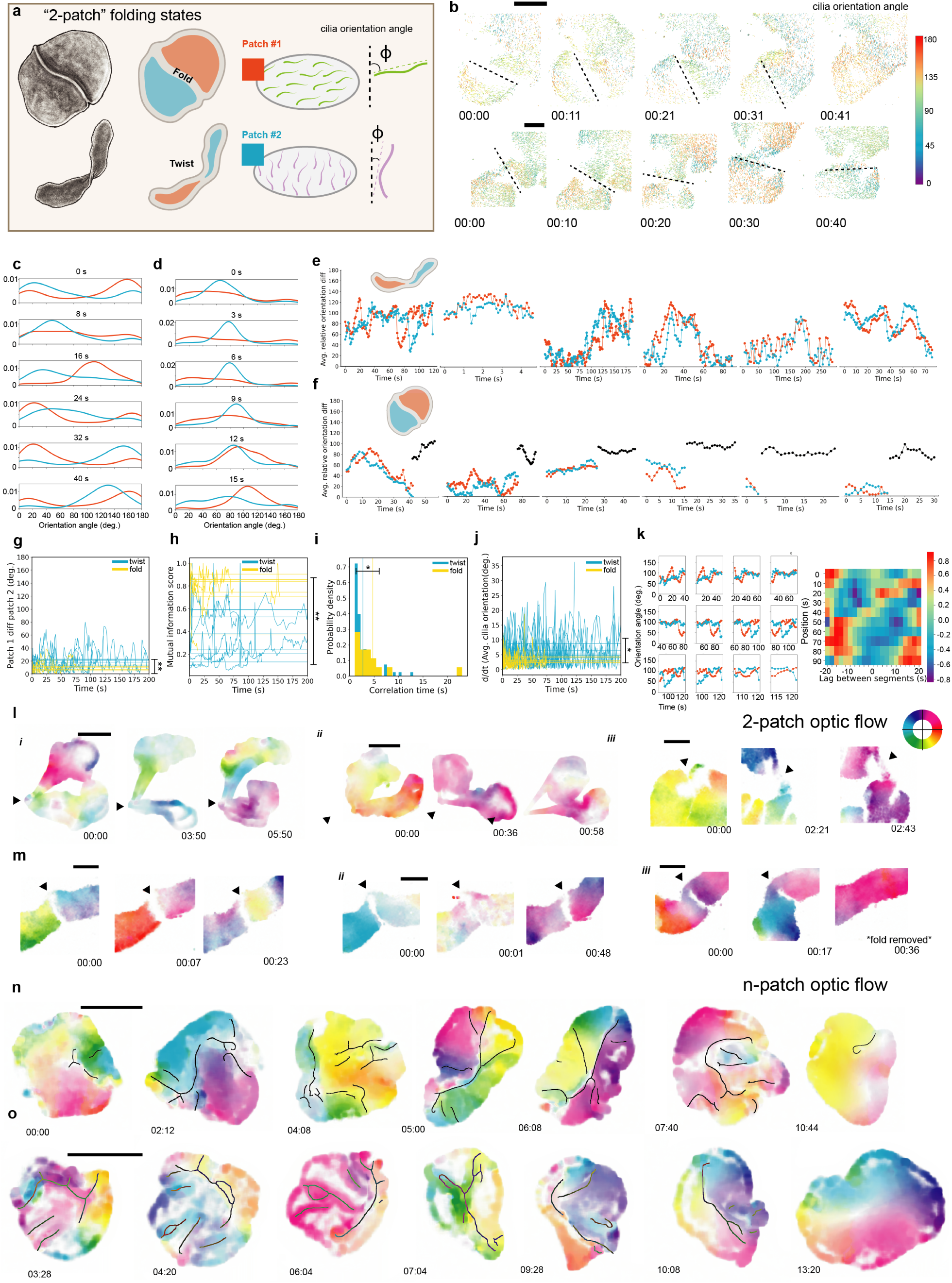
Cilia-resolved imaging and optic flow analysis of two-patch folded and twisted animals. (a) To study the impact of folds and twists on ciliary flocking behavior, we imaged simple “two-patch” folding configurations containing either a single fold or 2*π* twist and 2 flat patches (shown here in orange and blue), at single-cilia resolution. We then extracted cilia orientations across the body (see Methods). Previous work has established cilia flocking behavior in *T. adhaerens* wherein the orientations of neighboring cilia become correlated via elastic coupling [31]. (b) Example two-patch unfolding trajectories containing a single fold (top row) and single twist(bottom row) with extracted cilia orientations. The position and angle of the fold or twist is indicated by the dashed black line. Individual cilia orientation angles are indicated by color. These data enable the simultaneous visualization of ciliary dynamics and folding behavior. (c-d) Example patch-wise cilia orientation distributions of a folded (right) and twisted (left) animal. In both cases, the lack of coordination between patches is apparent; patches exhibit distinct peaks of cilia orientations angles. (e-f) Average patch-wise cilia orientation for six (e) twisted and (f) folded animals. In these time series, regions of both inter-patch correlation and decorrelation are apparent. Cilia exhibit a wide range of orientations relative to the twist or fold, from 0° to 140° for twists and 0° to 90° for folds. In (f), the black series represents the average ciliary orientation of the animal after the fold is removed. (g) Inter-patch orientation diff (difference between average ciliary orientations in each patch) for folded animals (yellow) and twisted animals (blue). Twisted animals exhibit a greater mean diff (horizontal lines) as compared to folded animals (*t* (10) = −4.39, p=0.0013). All animals exhibit inter-patch diff fluctuations in time. (h) Mutual information score between cilia orientation distributions for folded animals (yellow) and twisted animals (blue). Folded animals exhibit higher mean mutual information scores than twisted animals (*t* (11) = 4.38, p= 0.0011). (i) Patch correlation time distribution for all animals, defined as the length of time two patches continuously exhibit an average cilia orientation within 10° of one another. Folded animals exhibit longer correlations times as compared to twisted animals (Wilcoxon–Mann–Whitney test, p=0.023). Vertical lines indicate mean correlation times for each distribution. (j) Time derivative of average cilia orientation by patch. On average, patches in twisted animals reorient faster than those in folded animals (*t* (10) = −3.04, p=0.012). (k) Windowed time lag cross correlation of average cilia orientation for two patches. Introducing a time lag between patches leads to strong positive and negative correlations between patches. For example, correcting a 15 second lag between patch one and two from 50-80 seconds leads to a strong positive correlation between the average orientations of the two patches. (l-m) Optic flow between successive frames (separated by 2 seconds) of two-patch unfolding time lapses for (l) twisted animals and (m) folded animals. Color encodes the direction of tissue motion, which is correlated with ciliary orientations [31]. The orientation color map is shown in (l) (top, right). For both twists and folds, we observe cases of decorrelated patch motion (l.i, m.i) and correlated patch motion (l.ii, m.ii). In twisted animals, we also observe patch correlation arising from inter-patch contact (l.iii). In folded animals, we observe the transition from decorrelation in the presence of fold to correlation immediately following the removal of the fold (m.iii). (n-o) Optic flow between successive frames of two representative crease datasets from figure 3. Strikingly, patch motion is aligned with crease defect lines. In this way, creases act as information bottlenecks in cilia flocks. Scale bars are: (b, l.iii, m) 100 *µ*m and (l.i-ii, n,o) 500 *µ*m.

**Fig. 6:**
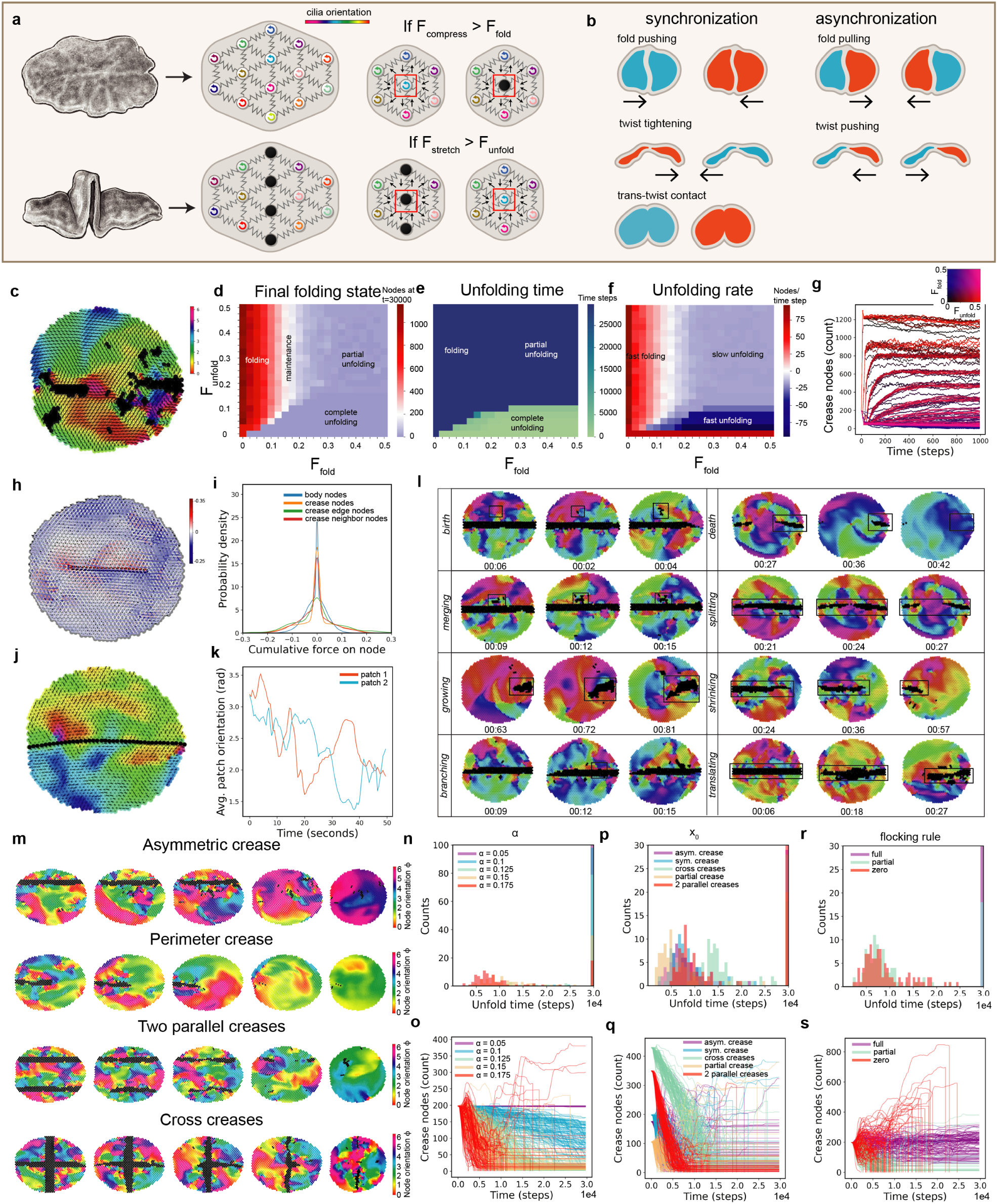
2D model of unfolding driven by cilia flocking. (a) Schematic of 2D active elastic cilia flocking model of folding and unfolding behavior. In this model, we represent folds as regions of inactive nodes (shown in black). Inactive nodes act as partial information barriers; they can pass cilia orientation information via compressive but not tensile forces, as shown experimentally in SI Movie 14. Parameters *F_fold_* and *F_unfold_* establish the thresholds for folding and unfolding. See SI for model description. See SI Movie 17. (b) Summary of inter-patch communication rules. The differing geometry of folds and twists leads to partial communication schemes wherein inter-patch communication in folds is enabled by compressive but not tensile forces. (c) Snapshot of unfolding simulation showing regions of active, flocking cilia and inactive creases. Color represents activity orientation, shown also as black lines associated with each node. (d-g) Effects of varying *F_fold_* and *F_unfold_* on unfolding behavior. Phase space of *F_fold_* and *F_unfold_* showing the (d) final folding state (number of crease nodes in the network at t=30,000 time steps), (e) unfolding time, and (f) unfolding rate (slope of crease nodes vs. time for the first 15 time steps of the simulation). The low *F_unfold_*, high *F_fold_* regime yields rapid, complete unfolding behavior. (g) Unfolding trajectories for all simulations shown in (d-f). A 2D color code (top, right inset) indicates the values of *F_unfold_* and *F_fold_*. (h) Snapshot of inter-node forces in a simulation with a partial crease (partial information bottleneck). Red indicates stretch and blue indicates compression between nodes. Qualitatively, we can observe the alignment of extreme inter-node forces with the central horizontal crease. (i) Probability density of inter-node force distribution for 30,000 time steps without any folding or unfolding permitted. Crease edge nodes, crease nodes, and crease neighbor nodes exhibit a higher probability to experience extreme stretch and compression as compared to body nodes. (j) Snapshot of cilia orientations with a full-body horizontal crease (information bottleneck). Qualitatively, we can observe distinct patches arising on either side of the crease, which shows the effect of the crease on neighboring cilia. (k) Time series of average cilia orientation in simulation with initial condition as shown in (j) for 30,000 time steps without any folding or unfolding permitted. As in Figure 5, we observe both correlation and decorrelation between the two patches, and clear cases of time-lagged correlation. (l) Unit operations of creases in our simulation. As observed in Figure 3k, here we also observe cases of crease birth, death, merging, splitting, growing, shrinking, branching, and translating arising from active node dynamics. (m) Representative simulations for varied initial folding states. (n-o) Impact of activity on unfolding behavior. When activity is too low, unfolding cannot occur. As activity is increased, slow and then fast unfolding occurs. For higher activity values, folding can also occur in some cases. Notably, when unfolding is observed, crease removal is non-monotonic; activity creates new creases while removing other creases. (p-q) Impact of initial condition on unfolding behavior. Partial crease unfolding is faster, while cross unfolding is slower. (r,s) Impact of inter-patch communication mode on unfolding trajectory and time. Partial and zero communication leads to fast unfolding while full communication does not enable unfolding.

### 2.5 Cilia-resolved imaging and optic flow analysis of two-patch folded and twisted animals

Our previous work in *T. adhaerens* has established that the microscale physics of individual walking cilia leads to large-scale emergent flocking behavior in the ventral ciliary carpet [31, 51]. This behavior is believed to be mediated by local elastic coupling between single cilia and their neighbors, leading to the correlated orientations of nearby cilia [31]. Thus, we hypothesized that folding motifs, which by definition break local elastic coupling, would limit flocking between adjacent patches of cilia.

To test this hypothesis, we performed single cilia-resolved DIC imaging (see Methods) [51] of animals in “two-patch” configurations, containing either a single fold or 2*π* twist (Fig. 5a). We then extracted the orientations of individual cilia [51] and binned them by patch identity (Fig. 5a-b, SI Movie 14). These data enabled us to compare the evolution of cilia orientations in each patch as the folding state evolves (Fig. 5c-f). From average and distribution data, it is apparent that there are cases of alternating inter-patch correlation and decorrelation in both fold and twist datasets (Fig. 5e-f). Furthermore, in folded animals, we observe the return to correlation immediately following the removal of the fold (Fig. 5b, 5m.iii). We quantified these dynamics by computing the inter-patch average orientation difference (ΔΦ̅) (Fig. 5g) and the mutual information score (Fig. 5h), both of which exhibit fluctuations over time. We also extracted correlation time distributions (defined as the length of time two patches spend with an average difference of less than 10 degrees), and the gradient of average patch orientation (Fig. 5i-j). These metrics interestingly show marked differences between fold and twist, which hints at the role of motif geometry in shaping ciliary behavior (Fig. 5g-j). We additionally performed optic flow analysis of folded and twisted animals, which revealed the tissue displacement field of the animal over time (previously shown to correlate with the cilia orientation field [31]) (see Methods). In these data, we also observed alternating states of correlation and decorrelation between the two patches (Fig. 5l-m). Taken together, these observations indicate that the presence of a fold or twist disrupts the elastic interaction network of the ventral cilia.

Next, we considered how folds and twists differ in their ability to shape behavior of the cilia. Interestingly, we observed instances of time-lagged correlation in both folded and twisted animals suggestive of delayed interaction between cilia patches (Fig. 5k). Given the distinct geometries of folds and twists, it is elucidating to consider how a given fold or twist impacts force transmission (Fig. 6b). A bulk force from a patch may either be compressive or tensile (toward or away from the fold/twist, respectively). Because of the structure of a fold and low bending modulus of the animal, a tensile force exerted by one patch in the direction of the fold should not be able to interact with the cilia in the other patch, as this tension will first be converted into fold removal. On the other hand, a tensile force exerted by one patch through a twist should lead to twist tightening and subsequent interaction with the other patch. Conversely, a compressive force on a fold should enable interaction with the neighboring patch, while a compressive force on a twist should lead to twist loosening and prevent interaction with the neighboring patch (Fig. 6b, SI Movie 15). In this way, a fold or twist effectively acts as an “information bottleneck” through which cilia orientation information (spin wave) from one patch is transmitted to the other patch [72]. This transmission may lead to a reciprocal or non-reciprocal interaction, depending on the elastic structure of the motif (i.e. fold or twist). This is a novel mode of communication within ciliary flocks, and flocks more generally, which has not been previously described.

We also considered how cilia are impacted by partial information bottlenecks in the context of perimeter folds (as defined in Fig. 2f). Because perimeter folds form creases with an exposed tip (around which cilia can pass orientation information), it is not apparent how they should impact the surrounding cilia flock. To address this question, we examined the cilia orientation patterns at the edge of perimeter crease defects, and observed the presence of distinct pinwheel orientation patterns visible in both DIC and optic flow data (SI Movie 16). Thus, even partial folds can still block paths of information flow, leading to defect lines in the cilia flock. These patterns may play a role in the crease merging dynamics highlighted in Figure 3 (SI Movie 13).

Finally, to test whether these phenomena can be generalized to more complex, *n*-patch systems, we performed optic flow analysis on the ventral crease time lapse data presented in Figure 3, and observe disjointed patches arising along crease lines (Fig. 5n-o). In other words, ciliary orientations show a dramatic jump across crease boundaries, thus exhibiting “patchy” flocking dynamics. This scheme leads to the establishment of quasi-compartmentalized, disjointed ciliary patches that can interact through bulk forces (Fig. 5n-o). Importantly, some degree of anti-correlation between ciliary patches is necessary for fold removal or twist tightening. These datasets demonstrate a remarkable “two-way coupling” mechanism wherein folds and twists generate ciliary flocking states which promote their own removal from the sheet via disjointed motion.

### 2.6 2D model of unfolding driven by cilia flocking

To further investigate the proposed two-way coupling between ciliary dynamics and folding state, we next modeled cilia-driven unfolding in a two-dimensional context. We build upon the active elastic resonator model for cilia flocking embedded in an epithelial sheet as an active solid (presented in Bull et al. [51], see SI for model summary). In order to capture the unique dynamics of unfolding behavior, we integrate out-of-plane folds into this flocking model as sources and sinks of active nodes (i.e., individual ciliated cells) (Fig. 6a). Inactive nodes represent cells within a fold, whose cilia cannot participate in flocking behavior. Active nodes may recruit inactive nodes into their patch (i.e., unfolding) by exerting tensile forces above a threshold, *F_unfold_* (function of *E_bend_* and *E_cc_* discussed in section 2.4). Conversely, active nodes may locally initiate inactive nodes (i.e., folding) via compressive forces above a threshold, *F_fold_* (function of *E_bend_* and *E_cs_* discussed in section 2.4). Active nodes move at each time step based on their activity *α*, and tuning this parameter relative to *F_unfold_* and *F_fold_* influences unfolding behavior (Fig. 6n-o). Figure 6c shows an example snapshot from an unfolding simulation using this model.

To determine the possible regimes of behavior in this system, we performed a parameter sweep of *F_unfold_* and *F_fold_* for a fixed activity value *α* (Fig. 6d-g). From the resulting phase spaces (Fig. 6d-f), we can predict the following behavioral regimes: 1) high *F_unfold_* and low *F_fold_* exhibits rapid, partial or complete folding, 2) high *F_unfold_* and high *F_fold_* exhibits slow, partial or complete unfolding, 3) low *F_unfold_* and low *F_fold_* exhibits rapid, folding or unfolding as a function of *F_unfold_/F_fold_*, and 4) low *F_unfold_* and high *F_fold_* exhibits rapid, complete unfolding. Given that *T. adhaerens* exhibits robust unfolding, we expect it to fall in the low *F_unfold_*, high *F_fold_* regime (see also section 2.4).

To test our intuition that information bottlenecks in flocking cilia can lead to their own removal, we ran simulations with very high *F_fold_* and *F_unfold_* values to inhibit any unfolding or folding behavior. This enabled us to quantify the distribution of compressive and tensile forces over time as a function of node type (e.g. active vs. crease nodes). Qualitatively, we observe the alignment of inter-node forces along the length of a crease (Fig. 6h, SI Movie 16). To quantify this observation, we obtained the distribution of inter-node forces over time, which revealed that crease nodes, crease edge nodes, and crease neighbor nodes have a higher probability of experiencing extreme stretch or compression as compared to body nodes (Fig. 6i). Interestingly, crease edge nodes exhibit the highest probability of experiencing extreme forces, which may be explained by their geometry and exposure to a greater number of neighbor nodes as compared to nodes within the crease interior. We also extracted the average patch orientation for simulations without any unfolding or folding, and found similar dynamics to those observed in Figure 5f. These results reinforce our intuition that crease defects both impact surrounding ciliary dynamics and promote their own removal through accumulation of ciliary forces (i.e. two-way coupling).

To further characterize unfolding dynamics, we next ran 100 simulations with low *F_unfold_* (0.1575) and high *F_fold_* (0.25) (SI Movie 17) in a background of fixed *α*. We then examined the resulting unfolding trajectories and identified common crease unit operations. Interestingly, we observe equivalents of the crease operations identified in real data in Figure 3k, which highlights the tendency for ciliary flocks to exhibit 2D spin waves whose fronts lend themselves to linear crease formation (Fig. 6l). We also considered the effect of initial folding condition on unfolding time (Fig. 6m, p-q). In experiments, it is difficult to replicate an initial condition, as we rely on the animal’s random settling to a substrate. Thus, a simulation approach allows us to ask how initial condition affects unfolding time. We tested five different initial conditions: symmetric crease, partial crease, asymmetric crease, cross creases, and two parallel creases (Fig. 6m). We find that unfolding can occur in all cases, however certain conditions led to longer unfolding times (Fig. 6p-q). For example, the partial crease condition led to the fastest unfolding times, while the cross crease condition led to the slowest unfolding times (Fig. 6p-q). This may be due to a combination of the total number of crease nodes and the initial geometry of the active patches relative to the crease positions. Interestingly, the cross creases are resolved one branch at a time (Fig. 6m), which hints at the importance of node compartmentalization in setting the unfolding time and trajectory.

Finally, we considered the role of inter-patch communication in driving crease removal (i.e. the strength of the information bottleneck). As discussed in section 2.5, folds act as information bottlenecks, which block direct flow of ciliary orientation information (i.e., spin waves) but allow partial information to leak via bulk forces (Fig. 6b, SI Movie 14). To test whether this partial communication scheme provides a benefit to unfolding behavior, we simulated unfolding with different inter-patch communication rules (Fig. 6r-s): full communication across creases (via both tension and compression, equivalent to no crease), partial communication (via compression only), and zero communication (complete blockage). Surprisingly, we find that the full communication mode leads to very slow unfolding times, while partial and zero communication enables fast unfolding (Fig. 6r-s). Thus, the inter-patch communication mode is critical for defining the unfolding rate and unfolding trajectories. It is crucial for active patches to achieve decorrelated motion to enable unfolding. Information bottlenecks within the flock enable this decorrelated motion, whereas full communication leads to the maintenance of folded states.

## 3. Discussion

Here we characterize folding and unfolding behavior in aneural animal *T. adhaerens* and discover a novel mechanism for behavioral tissue unfolding which is driven by ciliary flocks shaped by information bottlenecks (Fig. 7). In doing so, we establish a uniquely tractable living system for studying the mechanics of non-stereotypical thin sheet folding driven by distributed activity. Through experiments and simulation, we discover a remarkable two-way coupling in which folds and twists act as topological defects which break ciliary coordination, leading to the emergence of flocking states that explicitly remove these defects. In this way, *T. adhaerens* exhibits “embodied intelligence” for unfolding; it is mechanically suited for unfolding or folding, depending on its local substrate geometry. We believe the use of folding to generate information bottlenecks in cellular activity may be a more general feature of multicellular systems and invites further study, especially in the context of animal morphogenesis.

**Fig. 7:**
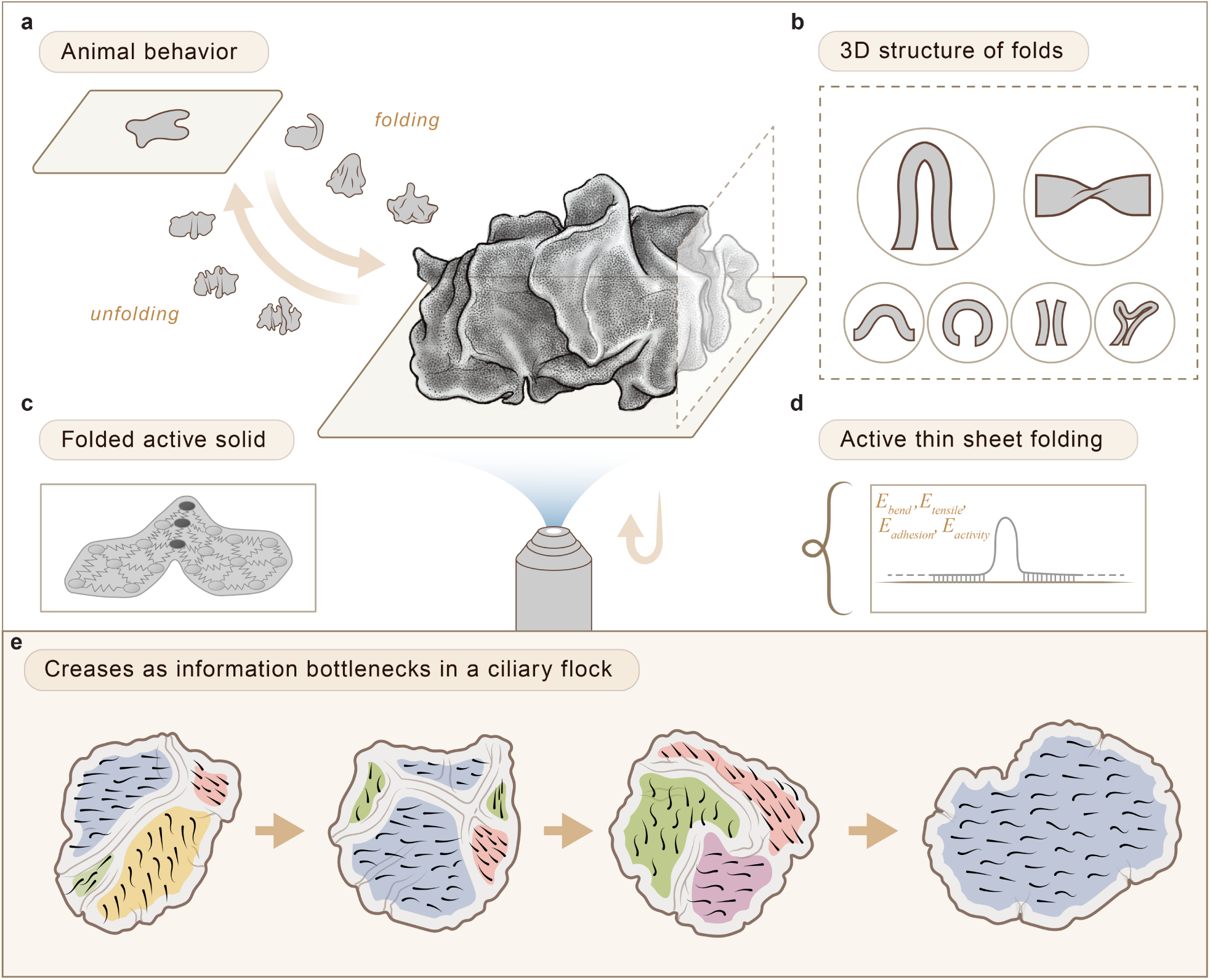
Graphical summary. (a) In this study we discover a characteristic, non-stereotypical thin sheet folding behavior exhibited by aneural animal *Trichoplax adhaerens*. This cilia-driven behavior enables the animal to transition between folded and unfolded states. In shallow ocean waters where *T. adhaerens* has been collected, this unfolding behavior is likely crucial for survival in an environment where patchy, turbulent flows can lead to surface detachment. (b) We study the folded state of the animal and identify 3D structural motifs such as rucks and hairpin folds. (c-d) Given these motifs, we model unfolding behavior using a 1D toy model of active unfolding and a 2D active solids model of unfolding driven by ciliary flocking. Through these approaches, we show the importance of ciliary activity in driving unfolding behavior; the impact of ciliary coordination schemes on unfolding efficiency; and the interplay between ciliary flocking dynamics and folding state. (e) We collect time lapse datasets of the animal’s active domain (the ciliated ventral surface in contact with the substrate) over the course of unfolding behavior. From these data, we discover an elegant mechanism for unfolding in which creases act as defects that break correlation in the ventral cilia flock. This broken correlation leads to the emergence of quasi-compartmentalized flocks which are capable of the disjointed motion required to remove folds. This “two-way coupling” between ciliary flocking dynamics and folding state is a remarkable example of embodied intelligence in a brainless animal and represents a powerful new model system in which to study active folding of thin multicellular sheets.

Our work primarily focuses on the role of mechanics in this physical folding phenomenon. Future studies will examine the relationship between biochemical signals and sheet mechanics in *T. adhaerens*. For example, it is known that neuropeptides can trigger folding in *T. adhaerens* [33], but the interplay between neuropeptide response and modulation of ciliary activity to trigger folding remains to be understood. Additionally, the high curvatures observed in folds (Fig. 2d) raise the question of cellular curvature sensing within *T. adhaerens*. It has been shown that epithelial sheets exhibit fast response to curvature [73], but few studies have been done to test the response to dynamically modulating curvature [74]. We have shown that *T. adhaerens* is a powerful model system for studying dynamic epithelial curvature response.

Additionally, the work presented here provides new perspective for the evolutionary history of multicellular folding. We demonstrate that ciliary activity (coupled to a rigid substrate boundary condition) is sufficient to drive tissue folding and unfolding without genetic control. Other examples of dynamic, mechanical folding have been recently noted in developmental systems [75]. It is likely that even before the evolution of molecular-driven epithelial tissue folding processes, like those classically studied in embryonic development, a multitude of other active cellular mechanisms (e.g. ciliary activity, flagellar beating, substrate adhesion, contractility, etc.) enabled tissue folding, perhaps allowing primitive multicellular organisms to achieve dynamic three-dimensional forms [16].

Beyond biology, our discovery of two-way couplings between fold/twist information bottlenecks and 2D flocking dynamics in an elastic sheet introduces a new wrinkle into active matter research. Active matter systems are known to exhibit topological defects which lead to global deformations of a material [76]. To the best of our knowledge, our work constitutes a novel class of non-conservative topological defects in an active flocking system. Unlike previously characterized topological defects, folding defects also act as material sources and sinks, and therefore have the power to determine the active members of a ciliary flock. The principle of topological information bottlenecks in flocks is general [72] and could potentially be engineered in other systems as a means to structure information flow. The ciliary flocking dynamics revealed here raise questions about a larger space of emergent collective behaviors made possible when the two-way coupling of topological defects and flocking dynamics is introduced.

Finally, we expect this discovery of living, multicellular active origami to inspire a new class of engineered living and non-living active origami processes, which rely on local, distributed activity to drive surface-attached unfolding. Active origami has been harnessed by engineers to build a variety of self-folding structures, including telescope sunshields [77], soft robots [78, 79], and micro-scale devices [80]. Our work makes significant progress toward understanding the programmability of biological self-folding processes.

## 4. Methods

### 4.1 Animal culture

Laboratory cultures of *Trichoplax adhaerens* (gift from L. Buss, Yale University) were maintained in 14 cm glass petri dishes of 100mL artificial seawater (ASW) at 19*^o^*C with an 18h light and 6h dark cycle. ASW was prepared by dissolving Coral Pro Salt Mix in Millipore water with a final concentration of 45 PPT. *T. adhaerens* cultures were fed weekly with 10 mL of concentrated *Rhodomonas lens* cultures. *R. lens* cultures were also maintained at 19*^o^*C with an 18h light and 6h dark cycle and provided with Guillard’s F/2 media (Micro Algae Grow, Florida Aqua Farms). On a weekly basis, existing culture dishes were refreshed and new culture dishes were prepared. Existing culture dishes were refreshed by replacing 1/3 the volume with fresh ASW (with 250*µl* Micro Algae Grow per liter) and 5-10 mL *R. lens* secondary culture. New dishes were prepared by adding 90 mL fresh ASW (with 250*µl* Micro Algae Grow per liter) and 10 mL *R. lens* secondary culture, waiting 2 days, and adding 20 animals from an older dish to the new dish. *R. lens* cultures were split on a weekly basis. 2/3 culture volume was replaced with fresh ASW and Micro Algae Grow (500*µl* per liter of ASW). The *R. lens* culture flasks were stirred for 1 minute every hour, and aerated with an air pump.

### 4.2 Unfolding trials

Animals were isolated from culture dishes using a p200 micropipette and transferred into a 5 mL glass bottom dish of ASW. Animals were left to wash (i.e. shed any attached algae) for 2 hours before all unfolding trials unless otherwise specified. All unfolding trials were carried out in glass bottom dishes cleaned with Millipore water and containing 5 mL ASW unless otherwise specified. To start an unfolding trial, a p200 micropipette was used to peel the cleaned animal from the glass substrate and transfer it into the imaging dish. For repeated trials, new dishes were used to avoid any effect of debris or material left behind by the previous animal. To make the initial condition of an unfolding trial as consistent as possible, each newly used animal was put through an unfolding trial before the subsequent recorded trials.

### 4.3 Unfolding trials on capillaries and microspheres

Animals were held in close contact to a glass capillary or microsphere using a micropipette until partial attachment to the surface had occurred. At this point, the pipette was removed leaving the animal to continue attachment to the substrate. 120 *µm* borosilicate glass microspheres and 170 *µ*m borosilicate glass capillaries were used.

### 4.4 Vertical tracking microscopy

The ‘Gravity Machine’ imaging system [44] was used to obtain datasets of *T. adhaerens’* folding state in the absence of a rigid substrate. Animals were loaded into the annular imaging chamber with 80mL ASW. ‘Squid’ software [81] relying on DaSiamRPN [82] was used to track suspended, sedimenting animals.

### 4.5 *In situ* observation of animals on spherical algae clusters

In laboratory cultures, *T. adhaerens* was observed attaching to and wrapping around spherical algae colonies which naturally occurred in our culture dishes (likely in co-culture with *Rhodomonas lens* cultures). A micropipette was used to gently transfer animal-algae complexes into imaging chambers for light sheet imaging (see section 4.11). Complexes were incubated in 1X CellMask Orange Plasma Membrane Stain in ASW for 10 minutes and washed in fresh ASW before imaging. See section 4.11 for light sheet imaging protocol.

### 4.6 Unfolding time measurements

Unfolding trials were performed as described in section 4.2 for 163 trials across 45 animals in total. Unfolding times were measured from the moment the detached animal makes contact with the glass substrate, to the time when all folds are removed from the animal. Due to variations in animal shape and size, some animals exhibit more persistent perimeter folds than others. In these cases, the ‘unfolding clock’ was stopped when the fold content plateaued and folds were only observed at the perimeter.

### 4.7 Size-dependence unfolding time measurements

Unfolding trials were performed and times extracted as described in the sections 4.2 and 4.6. Five small (*∼* 0.1 *mm*^2^ diameter) and five large (*∼* 1 *mm*^2^ diameter) animals were selected to undergo ten repeated unfolding trials each. Imaging was performed using a Nikon AZ100 microscope and bright field illumination. Animal size was extracted in Fiji. Linear regression was performed using the scikit learn linear regression python library.

### 4.8 2D tracking microscopy

The ‘Squid’ imaging system [81] was used for automated tracking and imaging of *T. adhaerens* unfolding time lapses.

### 4.9 Ciliary inhibition with LiCl and Calcium-free sea water

Unfolding trials as described in section 4.2 were performed across 10 animals. One at a time, animals were first made to unfold in ASW (pre-treatment condition), and then immediately transferred to a second dish containing either 100 mM LiCl in ASW or Calcium-free seawater (CFSW) for a second unfolding trial (treatment condition, left for 15 minutes). 100 mM LiCl solution was made by preparing a 2M stock solution in ASW (4.23 g LiCl into 50 mL ASW) and diluting 1:20 in ASW. Calcium-free seawater was prepared as described in [83] supplemented with EGTA (2mM final). After this step, animals were transferred to a fresh dish of ASW for a 10 minute recovery, after which they were transferred to a fourth dish of ASW for a final unfolding trial (post-treatment condition). Time lapses were collected using ‘Squid’ [81]. Animal contact area contours were extracted in Fiji.

### 4.10 Manual tracking microscopy

For live unfolding data collected on the Nikon AZ100 (Fig. 1j-k, 1n-p), Bruker Luxendo TruLiv3D Imager (Fig. 1b, Fig. 2), or Nikon TI2 (Fig. 3, Fig. 5), manual tracking was used to keep the animal within the field of view throughout the unfolding time lapse.

### 4.11 Light sheet microscopy

Light sheet imaging of *T. adhaerens* was done using the Bruker Luxendo TruLive3D Imager at Nikon CFI APO LWD 25x 1.10 NA objective and 100 mm tube lens to achieve a 12.5x magnification. Fluorinated ethylene propylene (FEP) foil (rather than glass) chambers were used, as required for this microscope. The minimum exposure time allowed by the microscope (11 ms) was used per z-slice in order to minimize blur in live samples. Both CellMask Plasma Membrane Stain and LysoTracker were separately used to generate a strong fluorescent signal throughout the animal’s entire body, which enabled visualization of the folding state. 3D data processing and analysis (including surface fitting) was done in Imaris and 2D data processing and analysis was done in Python.

### 4.12 Curvature calculations

Local Gaussian curvature of 3D surfaces was extracted in Meshlab using the compute curvature principal directions scale dependent quadratic fitting method. Local curvature of 2D cross sections was extracted in Python using a window size of 10 pixels and calculating local curvature as *|d*^2^*y/dx*^2^*|/*(1+(*dy/dx*)^2^)^3^*^/^*^2^.

### 4.13 Chemical fixation

Animals were isolated and washed in fresh ASW for 2 hours to remove any algae residue and stained as desired. After staining, animals were transferred into a fresh dish of ASW and triggered into an unfolding trial via peeling with a p200 micropipette. Folding state was monitored by a dissection scope. When sufficient ventral-substrate attachment had occurred, paraformaldehyde (PFA) was added to the dish to a final concentration of 2%. After fixation in PFA for 30 minutes, fixed animals were washed twice in phosphate-buffered saline (PBS) and imaged immediately.

### 4.14 Confocal microscopy

Animals were fixed as described in section 4.13 and imaged using the Zeiss LSM 780 multiphoton laser scanning confocal microscope with a Plan-Apochromat 63x/1.40 Oil DIC M27 objective. Animals were stained with Hoechst and CellMask Plasma Membrane stain prior to fixation to enable fluorescent visualization.

### 4.15 Perimeter crease length and area measurements

Across all unfolding datasets, nine events were identified in which a single perimeter crease increased or decreased in length without any other folding or unfolding events taking place in the animal’s body. From these events, contact area and crease length were extracted in Fiji. Total area within the fold was extracted by subtracting the animal contact area from the total animal area in the flattened state.

### 4.16 Crease imaging, segmentation, and analysis

Unfolding time lapses were collected as described in section 4.2 and imaged on a Nikon TI2 inverted microscope using a 10X 0.3 NA objective. To enable crease visualization and segmentation, fluoroscein was added to the imaging dish to a final concentration of 0.001%. Image processing was done in Fiji. Automatic thresholding combined with manual correction was used to segment crease lines and animal contours. Binary crease images were then skeletonized, and skeletons were used as an input to analysis pipelines. Feret diameter and roundness metrics were extracted in Fiji. Crease tracking was done in Fiji using the MTrack2 plugin. All other image analysis was done in Python using OpenCV. Unit operations were identified through manual search. Network analysis of ciliary domains was done by skeletonizing the negative space created by creases and counting the number of edges in the resulting skeletons.

### 4.17 Cilia imaging, segmentation, and analysis

Unfolding time lapses were collected as described in section 4.2 and imaged on a Nikon TI2 inverted microscope using a 60X 1.49 NA oil objective. Smaller animals were used in order to increase the likelihood of reaching a simple, two-patch state, and maximize the fraction of the entire animal visible in the field of view. Repeated unfolding trials were performed (as described in section 4.2) until the animal reached a two-patch state. The progression of the two-patch state was imaged and manually tracked. The z-plane was continuously adjusted to ensure cilia appeared as white lines, as necessary for segmentation. Individual cilia were segmented in Fiji using a bandpass filter to remove tissue background. Size and shape filtering were used to clean the image, and isolate binarized cilia. Binarized images were analyzed in Python. PCA was used to obtain the eigenvectors and subsequent orientation of each cilium contour [31]. Images were manually annotated with the patch identity and fold or twist orientation relative to the horizontal, from which cilia were binned into patches and orientations were reported relative to the fold or twist orientation. Mutual information score was calculated using scikit learn feature selection mutual info regression method.

### 4.18 Optic flow

Dense optic flow was calculated between successive frames of unfolding time lapses using the Gunnar Farneback’s algorithm implemented in OpenCV in Python.

### 4.19 1D scaling analysis

To perform scaling calculations, we used existing measurements for elastic properties of epithelial sheets and adhesive/active properties of cilia. Previous measurements of MDCK epithelia (which are a similar sheet thickness to *T. adhaerens*) yielded a bending modulus *B ∼* 2 *×* 10*^−^*^13^ *N · m* and Young’s modulus *Et ∼* 1.5 *×* 10*^−^*^2^ *N/m* [27]. Force spectroscopy assays of single-cell *Chlamydomonas* flagellar surface adhesion revealed a median adhesion force of 1.4 nN per flagellum [69]. Dynamic micropipette force measurements in *Chlamydomonas* revealed flagellar beating forces of *∼* 26 pN per flagellum [70] and magnetic micro-bead assays of individual human airway cilia revealed beating forces of *∼* 62 pN per cilium. We use the average value of these two measurements (44 pN per cilium). Finally, we measure the ciliary density on the ventral surface of *T. adhaerens* through direct ciliary imaging (as described in section 4.17) and counting the number of visible cilia in a *∼* 50 *×* 50 *µm* bounding box, which gave a ciliary density of 6 *×* 10^10^ cilia/m. Additionally, using ventral cell size data from Smith et al. [30], we obtain an estimated ciliary density of 2 *×* 10^11^ cilia/m. We use the average value of these two estimates (*ρ_c_* = 1.3 *×* 10^11^ cilia/m). From cilia imaging data, we can also estimate the average ciliary length to be *∼* 8 *µm*, and based on height fluctuation imaging in Bull et al., use *l_c_* =*∼* 3 *µm* (the distance over which ciliary adhesion force acts to bring the ventral epithelium closer to the substrate) [51]. These measurements allow us to perform energy estimates in *T. adhaerens*. See SI for further details.

### 4.20 1D simulations

1D cellular strings were simulated as a length *n* lattice representing *n* cells in contact with a rigid substrate. Using this framework, we focused specifically on the dynamics of hairpin folds and twists, as defined in section 2.2. In our model, *E_cc_* represents the adhesion energy between individual cilia in contact with one another within folds or twists. *E_cs_* represents the adhesion energy between a cilium and a substrate in flat, substrate-adherent regions of the string. *F_c_* is implemented as a virtual activity force exerted by each cell in flat regions, which can bias the motion of fold and twist boundaries to the left or right. In each time step of the simulation, we locally perturb the positions of fold and twist boundaries, and choose the lowest energy configuration option based on the influence of *E_cc_*, *E_cs_*, and *F_c_* across the sheet. 1D simulations were written in Matlab. See SI for further details.

### 4.21 2D simulations

2D active solids were simulated as active elastic spring networks where each node in the network represents a ciliated cell with a ciliary activity *α* and ciliary orientation *ϕ* [31]. In each simulation time step, node *α* and *ϕ* values initiate node displacements, which introduce torques on the virtual cilia of neighboring nodes, leading to the subsequent update of their *ϕ* values [31]. All unfolding simulations were run with the same initial folding state (a symmetrical two-patch state containing a single fold with an initial width of 5 nodes) unless otherwise specified; and a randomized initial *ϕ* field to mimic the initial state of contact after detachment and settling to a substrate. Simultaneously, inter-node forces throughout the sheet were tracked. Local compressive forces greater in magnitude than *F_fold_* lead to the initiation of inactive nodes which move with the sheet but cannot participate in flocking (i.e. folding). Tensile forces greater in magnitude than *F_unfold_* lead to the conversion of inactive nodes to active nodes (i.e. unfolding). 2D simulations were written in Python. See SI for further details.

## Acknowledgements

We thank all members of the Prakash Lab for helpful discussions. We especially thank Matthew Bull for discussions on cilia orientation extraction and for sharing code from previous 2D ciliary flocking simulations without creases. We thank Rebecca Konte for assistance with figure art and design. We thank Hongquan Li for SQUID-based tracking microscopes. We thank the Stanford Cell Science Imaging Facility. CMB acknowledges funding support from the NIH/NIGMS Cell and Molecular Biology Training Grant (5T32GM007276) and the NSF Graduate Research Fellowship Program (GRFP). MP acknowledges financial support from Schmidt Futures Innovation Fellowship, Moore Foundation, Dalio Foundation, NSF Center for Cellular Construction, NSF GCR Convergence Grant, and Woods Institute for the Environment.

## Author Contributions

CMB and MP designed the research. CMB collected the data. CMB analyzed the data in discussion with MP. CMB developed the 1D and 2D computational models in discussion with MP. CMB and MP wrote the manuscript.

