## Supplementary Information for "Creased ciliary flocks shape unfolding dynamics via information bottlenecks in an aneural animal"

#### Contents

**1. 1D Scaling Analysis** **2**

**2. 1D Toy Model Description** **4**

**3. 2D Cilia Flocking Model Description** **5**

**4. Perimeter fold length and area scaling** **8**

**5. Supplementary Figures** **12**

**6. Supplementary Movies** **18**

### 1. 1D Scaling Analysis

Here we consider the energetics of 1D active folding/unfolding. We begin with the schematic shown in Figure 1b. Comparing the associated energies allows us to understand where *T. adhaerens* lives in the phase space of active folding. For an ultra-thin epithelial sheet, cell-cell junctions prevent 2D flow in the tissue. Given the thickness of this animal $t \sim 20$ um, and typical length $L \sim 1$mm, these living sheets are high aspect ratio structures where $L/t \sim 50$. Since in-plane stiffness scales linearly to thickness (Young's modulus $Y \sim Et$) while bending modulus scales $B \sim Et^3$, we can easily see that the sheet can relax in-plane stress by bending out of plane (SI Movie 1, SI Movie 12). Moreover, as seen in three-dimensional light sheet and confocal datasets (Fig. 2d, 2f), these folds have a defined, crease-like geometry. For simplicity, we consider a 1D version of such a sheet where $\lambda$ is the crease width and $h$ is the crease height, with in-plane stress $\epsilon = \Delta/L$. Excess length that has been absorbed in the crease-like folds here is $\Delta$ (Fig 4b). Although it is difficult to directly measure the Young's modulus and bending modulus of a living animal with active cilia, we can reference past measurements of thin epithelial sheets of similar thickness ($t \sim 18$um) with in-plane Young's modulus being $Y \sim 10^{-2}$ $N/m$ and bending modulus being $B \sim 2.10^{-13}$ $N \cdot m$. We can further evaluate a Föppel-von Kármán number for our problem being $YL^2/B \sim 5 \times 10^4$ which is much larger than 1. Thus in a passive epithelial sheet of such a small thickness, bending out of plane is the preferred mechanism for release of any in-plane stress enabling dynamic crease formation (Fig. 3).

Next, we compare relevant energy scales in the problem. We have already looked at $E_{tensile}$ and $E_{bend}$ above for a thin elastic sheet. Two additional energies must be considered for an active sheet like *T. adhaerens*. As the name suggests, *T. adhaerens* utilizes cilia to adhere to any given substrate, allowing it to remain attached to a substrate geometry while still being able to move on the surface utilizing a unique ciliary walking gait (Fig. 1f-h) [1, 2]. This unique capability to both adhere to a substrate while still remaining motile arises simultaneously from ciliary dynamics in contact with a substrate [1, 2]. Thus, we introduce cilia-substrate adhesion ($E_{cila:substrate} \sim F_{cs}\rho_c\lambda$, later abbreviated as $E_{cs}$) where $F_{cs}$ refers to adhesion energy per unit cilium and $\rho_c$ refers to the number of cilia per unit length [3] and active tangential forces generated by the ciliary walking ($E_{activity}$). Based on the previous description of ciliary beating dynamics with a non-holonomic constraint, active forces can be calculated for a walking cilia [2]. Here we simplify this activity of walking cilia as $E_{activity} \sim F_c\rho_c Ln\Delta$, where $F_c$ is the tangential force generated per cilium.

Thus, the total energy of a creased organism on a given substrate at any given time $E_{total} \sim E_{cs} + E_{bend} + E_{tensile} + E_{activity}$. Unlike traditional passive folding problems, the system injects energy at the smallest length scale (ciliary walking) where a two-way coupling exists between geometry (crease folding patterns) and ciliary flocking (induced by activity). We can define a few non-dimensional energy scales including $\alpha = E_{bend}/E_{activity}$, $\beta = E_{tensile}/E_{activity}$, and $\gamma = E_{cs}/E_{activity}$. Comparing estimated values from experiments, we find that *T. adhaerens* lies in a regime where $\alpha < 1$, $\beta < 1$, and $\gamma \sim 1$. Thus, activity alone, in competition with adhesion at small sheet sizes, can induce crease folds and tensile strain in the system, leading to a rich dynamics of crease evolution, as seen in Figure 3. SI schematic 1 outlines the phase space of active unfolding in terms of $\alpha$, $\beta$, and $\gamma$. Another important consideration is to explore if gravity plays any role in these dynamics. To test this, we performed upside-down unfolding experiments and found that the animal is still capable of normal unfolding behavior (SI Fig. S10), implying the $E_{gravity} \sim \rho g t h$ is small compared to other energy scales in our problem.

In the past, activity in thin elastic sheets has only been explored either in context of thermalized sheets where activity appears as thermal noise enabling surprising stiffening of ultra thin films [4], or as growth in bacterial bio films that can buckle out of plane forming unique vascular patterns [5]. Unlike these two examples, surface activity enabled by flocking cilia [1, 2, 6] in *T. adhaerens* shows coherence over long length scales leading to a unique two-way coupling with emergent folding geometries and flocking patterns.

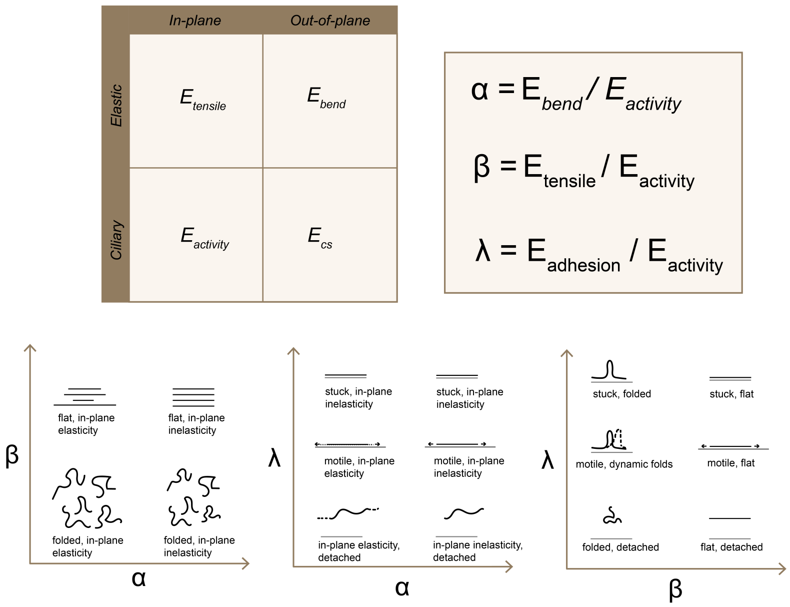


**SI Schematic 1**: **Energetics of active unfolding.**

#### 2. 1D Toy Model Description

Here we build a 1D toy model of unfolding behavior. We represent an unfolding animal as a 1D array of length $L$. The $L$ sites in the array are populated with id labels according to the current folding state. Based on the commonly observed folding motifs detailed in Fig. 2, and for simplicity, we focus on 3 possible folding motifs: 1) flat, surface attached (id=0), 2) $2\pi$twist (id $< 0$), 3) large-slope fold (id $> 0$). For example, $[\mathbf{0,0,0,0,1,1,1,1,1,0,0,0,0}]$ is an $L = 13$ sheet representing a single, central large-slope fold (id=1) flanked by two distinct flat regions (id=0) and $[\mathbf{0,0,0,0,0,0,-1,0,0,0,0,0,0}]$ is an $L = 13$ sheet representing a single, central twist (id=$-1$) flanked by two distinct flat regions (id=0). Flat regions in the sheet are assigned an activity direction right or left, and flat patches are fully coherent in their activity. For example, the corresponding activity array for the above folded sheet could be $[\mathbf{-1,-1,-1,-1,0,0,0,0,0,1,1,1,1}]$ where -1 indicates a leftward activity bias and 1 indicates a rightward activity bias. Activity here is a "virtual bias", rather than an explicit force. For example, if two patches flanking a central fold both exhibit rightward activity bias (and if activity strength is sufficiently high), then they will bias the motion of the fold to the right. However, if the two patches exerted the same rightward force, the fold would not move; rather the entire sheet would move to the right. Thus, activity implies forces rather than explicitly representing them; the two patches exhibiting rightward activity bias implies that the left patch is exerting greater force as compared to the right patch, leading to the translational motion of the fold to the right. This approach simplifies the model and enables the consideration of translational motion without requiring differential forces across patches. We assign the following parameters in the sheet: $E_{cs}$ for the adhesion energy between a single cilium and the substrate in flat regions; $E_{cc}$ representing the adhesion energy between 2 cilia within a fold; $F_c$ representing the active force of a walking cilium.

At each time step, we track the positions of the junction pairs defining the bounds of all folds and twists in the system ((4 left, 8 right) for fold id 1 and (6 left, 6 right) for twist id -1); locally perturb boundary junctions in all possible directions ((right,right), (right,left), (left,right), (left,left), (right,none), (left,none), (none,right), (none,left); and obtain the energy of the transition and the resulting state $i$ using the following calculation:

$$E_i = \alpha E_{cc} + \beta E_{cs} + E_{bend} + X(F_c) + v_{fold} + v_{twist} + \chi_{\text{twist}} + \chi_{\text{fold}}, \quad (1)$$

where $\alpha$ is the total number of sites in a fold or twist and $\beta$ is the total number of flat sites. We assume $E_{bend}$ to be negligible given the discussion in SI section 1 above. Function $X$ calculates how much energetic cost or benefit to provide a given state based on the orientation of flat patch activity and the directionality of the perturbation. $X(F_c) = (|lf|lc + |rf|rc)F_c$;, where left force $lf$ and right force $rf$ are equal to the number of flat sites in the patches to the left and right of the relevant motif, respectively, and left cost $lc$ and right cost $rc$ are equal to the sign of the activity of the left and right patches, respectively. $v_{fold}$ and $v_{twist}$ represent the cost of sliding regions of self-contact, are equal to the adhesion energy within the fold or twist, $\alpha E_{cc}$. This value is expected to be high for symmetric junction pair motion but low for asymmetric junction pair motion. Finally, $\chi_{\text{twist}}$ and $\chi_{\text{fold}}$ refer are used to assign infinite energy cost to topological rule-breaking within the simulation. $\chi_{\text{fold}}$ is always equal to zero, because all local junction pair perturbations are topologically acceptable. However, $\chi_{\text{twist}}$ is set to infinity for all asymmetric perturbations, which are not permitted by a twist (a twist may move to the left or right but any outward or inward boundary growth would require sheet rupture and reattachment, which is not permitted in this model). Furthermore, we do not consider the role of twist tightness here and assume a twist is always a point defect.

For sheets containing multiple folding motifs, the energies associated with all permutations of junction boundary motion are calculated and the lowest energy case selected. In the case of adjacent motifs, which are perturbed one at a time, conflicts are resolved by trying all possible combinations and selecting the lowest-energy configuration. The need to consider all permutations of local junction pair perturbation limits the total number of motifs in the sheet that the simulation can handle to 5-6, which is sufficient to explore the dynamics of patch activity and motif motion.

In summary, in this model, folds and twists are delimited by their boundary sites, the positions of which are locally perturbed left or right and moved at each time step according to the lowest energy configuration. Flat domains have directional (right, left) activity, which can bias the motion of motif boundaries. When activity $F_c$ is equal to zero, the system evolves according to the ratio of $E_{cc}$ (energy associated with cilia:cilia adhesion within folds) and $E_{cs}$ (energy associated with cilia:substrate adhesion along flat domains). When $E_{cc}/E_{cs} > 1$, the system evolves toward a flattened state. When $E_{cc}/E_{cs} < 1$, the system evolves toward a folded state. When activity is sufficiently high so as to dominate over adhesion energy, the system is capable of reaching a folded or unfolded state, depending on activity directionality over time.

### 3. 2D Cilia Flocking Model Description

Here we build upon the cilia flocking model previously described in [2] in order to study the interplay between ciliary flocking of flat patches and evolution of folds. Using this approach, we model *T. adhaerens* as an active elastic spring network. For a more complete discussion of the physics underlying ciliary walking and flocking in *T. adhaerens*, please see [1] and [2]. We create a hexagonal array of n nodes and with it initialize a sheet object defined by a node and crease list. Each node has the following associated variables at a given time: cilia orientation $\phi$, cilia height $\psi$, 2D coordinate $(x, y)$, and folding state $f$. The simulation parameters are listed and described below:

| **Parameter** | **Description** |
|---|---|
| $\Delta t$ | time step |
| $\sigma_{\text{pos}}$ | random noise in cell position update |
| $\sigma_{\psi}$ | random noise in cilia height update |
| $l$ | inter-node equilbrium distance |
| $\alpha$ | activity; ciliary force exerted on nodes to update positions |
| $\epsilon$ | driving amplitude |
| $\Gamma$ | reorientation timescale |
| $\Omega$ | natural frequency of ciliary beat |
| $J$ | ratio of timescales between phase change and tissue displacement |
| $F_{unfold}$ | force threshold for converting crease nodes to active nodes |
| $F_{fold}$ | force threshold for converting active nodes to crease nodes |

At each time step, the force exerted on each node $i$ by its $j$ neighboring nodes is calculated as the gradient of the elastic energy [2], which is given by:

$$F_{x,i} = \sum_j (x_i - x_j) * (\sqrt{(x_i - x_j)^2 + (y_i - y_j)^2} - l) \tag{2}$$

$$F_{y,i} = \sum_j (y_i - y_j) * (\sqrt{(x_i - x_j)^2 + (y_i - y_j)^2} - l) \tag{3}$$

where $l$ is the inter-node distance at equilibrium. These computed forces are then used to update the x,y coordinates of all nodes using the following equations:

$$\Delta x = \Delta t(F_x + \sigma_{\text{pos}} - \alpha(1 + \epsilon * \cos(\psi)) \cos(\phi)) \tag{4}$$

$$\Delta y = \Delta t(F_y + \sigma_{\text{pos}} - \alpha(1 + \epsilon * \cos(\psi)) \cos(\phi)) \tag{5}$$

The process of node position update (tissue motion) exerts a torque on cilia orientations:

$$\tau_{\mathrm{i}} = \cos(\phi_{\mathrm{i}})F_{y,i} - \sin(\phi_{\mathrm{i}})F_{x,i} \tag{6}$$

$$\phi_{\mathrm{i}} = \phi_{\mathrm{i}} - \Delta t \Gamma \tau_{\mathrm{i}} \pmod{2\pi} \tag{7}$$

Next, if including cilia height fluctuations, cilia heights are updated or otherwise kept at 1:

$$c_{\psi,i} = \sum_j \psi_{\mathrm{i}} - \psi_{\mathrm{j}} \tag{8}$$

$$\psi_{\mathrm{i}} = (\psi_{\mathrm{i}} + \Delta t(\Omega - Jc_{\psi,i} + \sigma_{\psi})) \pmod{2\pi} \tag{9}$$

Next, nodes must be checked for crease update. In the initial condition, nodes are manually prescribed as crease nodes or active nodes. Active nodes have $\alpha > 0$ while crease nodes have $\alpha = 0$. Crease node behavior is set according to the inter-patch communication rule: zero communication, partial communication, or full communication. With the partial communication rule, crease nodes may pass compressive forces only to neighboring active nodes, but not extensile forces. With the full communication rule, crease nodes may pass both compressive and extensile to neighboring active nodes. With the zero communication rule, crease nodes may not pass any forces to neighboring active nodes. In any case, active nodes exert forces on crease nodes, which enables crease nodes to move with the evolving sheet. To update creases at each time step, we must calculate extensile and compressive forces on all active and crease nodes, and compare these forces to folding and unfolding thresholds $F_{unfold}$ and $F_{fold}$, respectively. This is done by calculating the spring length at each time step as:

$$d_{i,j} = (\sqrt{(x_i - x_j)^2 + (y_i - y_j)^2} - l) \tag{10}$$

We then check for all nodes $i$ whether $d_{i,j} > F_{unfold}$ or $d_{i,j} < -1 * F_{fold}$. We then update the node identity accordingly for the next time step. For example, consider the following snapshot from a simulation and the highlighted 7-node neighborhood.

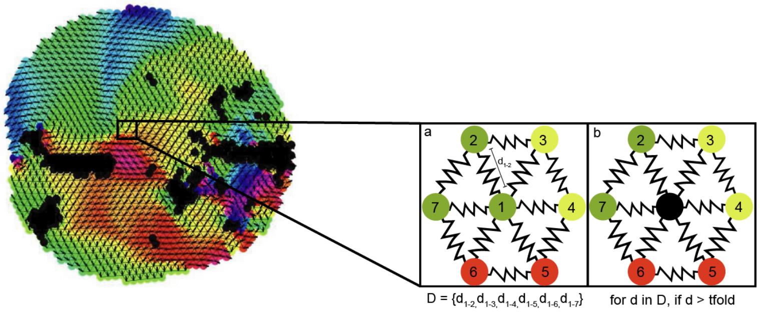


**SI Schematic 2**: **2D active elastic network unfolding model**

At time t, nodes 1-7 will each move a displacement $x_i$ in direction $\phi_i$, representing the cilia angle. These ciliary steps lead to changes in the relative distances $d_{1-2}, d_{1-3}, d_{1-4}, d_{1-5}, d_{1-6}, d_{1-7}$. If any of these distances $d_i > F_{fold}$, node 1 is converted from an active node to a crease node (panel b). At time $t + 1$, if $d_i > F_{unfold}$, node 1 is converted from a crease node to an active node (panel a).

### 4. Perimeter fold length and area scaling

Here we examine the scaling relationship between fold length and fold area. We consider a perimeter fold with height $h$, length $L$, fold angle $\theta$, and total surface area $a$ as depicted in the following schematic:

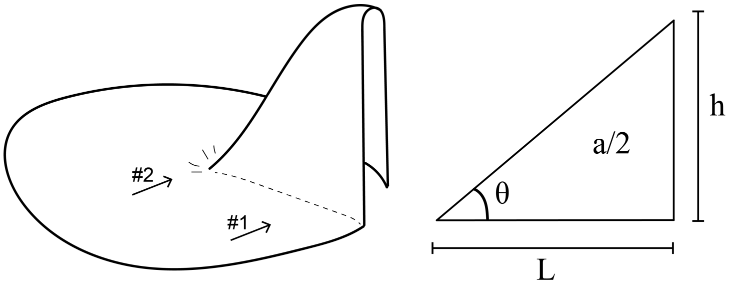


**SI Schematic 3**: **Perimeter fold dimensions**

Material can be added to or removed from this fold via local tissue action, as indicated by arrows 1 and 2 in SI Schematic 3. When new material is added to the fold, total fold area $a$ is increased and this can occur in one of three ways, as outlined in SI Schematic 4: 1) $h$ and $L$ must increase (constant $\theta$ case), 2) $\theta$ and $h$ must increase (constant $L$ case), or 3) $L$ must increase while $\theta$ decreases (constant $h$ case). Conversely, shrinking can occur along the same lines: 1) $h$ and $L$ must decrease (constant $\theta$ case), 2) $\theta$ and $h$ must decrease (constant $L$ case), or 3) $L$ must decrease while $\theta$ increases (constant $h$ case).

Interestingly, these processes can be likened to the stages of droplet growth on a surface. When volume is added to a droplet on a surface, constant $L$ growth will occur until the base of the droplet reaches a critical surface contact angle $\theta_a$. If further volume is added, the droplet will undergo constant $\theta$ growth. Conversely, if volume is removed from the droplet, it will undergo constant $L$ shrinking until reaching a critical angle $\theta_r$. If further volume is removed, the droplet will undergo constant $\theta$ shrinking. For droplets, these dynamics occur due to the interplay between liquid-liquid and liquid-surface interactions [7].

We can consider the analogous dynamics in folds. We focus on the case where material is added to the fold at the perimeter (i.e., arrow 1 in SI Schematic 3). When material is added at the edge, the fold will undergo constant $L$ growth until the fold reaches a critical angle $\theta_a$. Due to the presence of a d-cone-like singularity at this region (where the base of the fold meets the rest of the sheet), this critical angle should be a function of the elastic properties of the material. If further material is added, the fold will undergo constant $\theta$ growth (which we observe as fold lengthening). Conversely, if material is removed from the fold (again via tissue action at arrow 1), the fold will undergo constant $L$ shrinking until reaching a critical angle $\theta_r$. If further material is removed, the fold will undergo constant $\theta$ shrinking until flattening out. Notably, for both folds and droplets, constant $h$ growth is possible. However, in the case of folds, this requires tissue action at the tip of the fold (arrow 2 in SI Schematic 3).

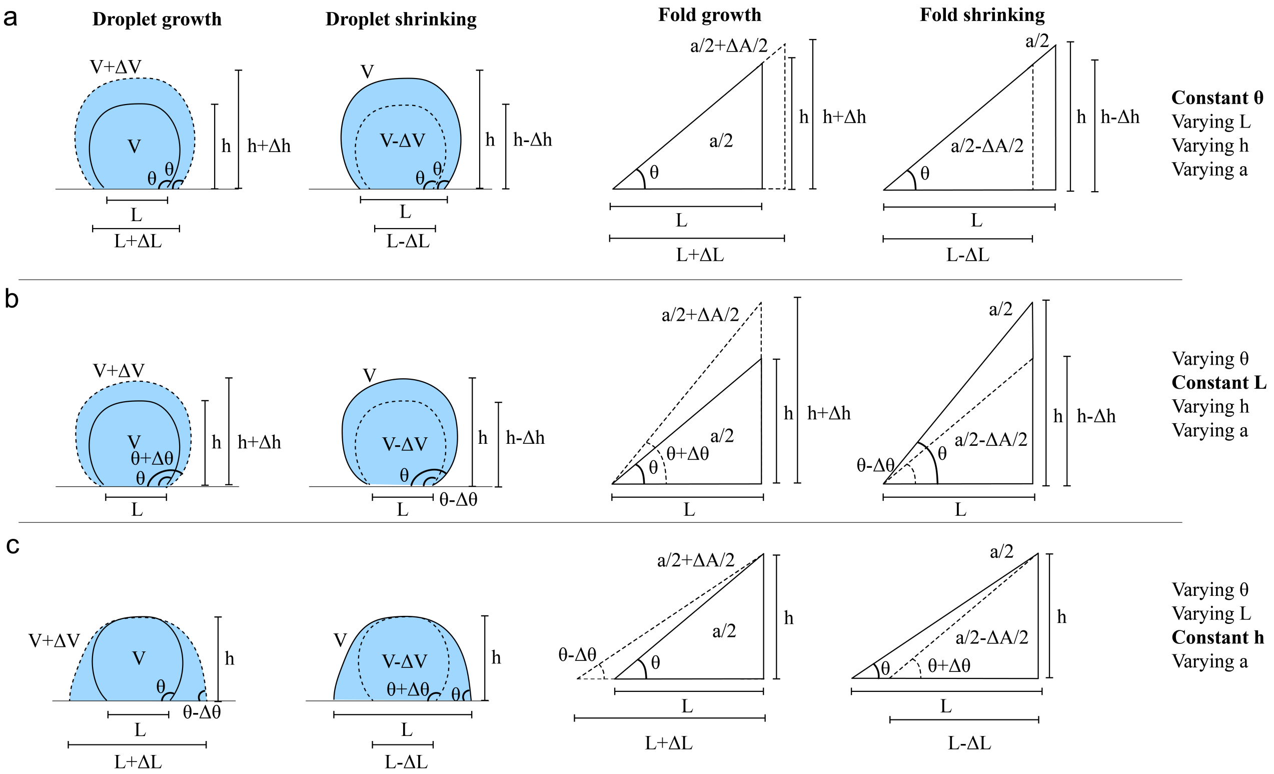


**SI Schematic 4: Growth and shrinking processes for droplets vs. perimeter folds**

With these growth and shrinking processes in mind, we use a simple geometric relation to estimate how total fold area scales with fold length in the case of constant $\theta$ and constant $h$ growth and shrinking.

#### Constant $\theta$ growth and shrinking

We first note the simple relationships:

$$L \tan\theta = h \tag{11}$$

$$(L + \Delta L) \tan\theta = h + \Delta h \tag{12}$$

We can write the equation for the area of the larger fold as:

$$\frac{a}{2} + \frac{\Delta A}{2} = \frac{1}{2}(L + \Delta L)^2 \tan\theta \tag{13}$$

Solving for $\Delta L$, we find that:

$$\Delta L = \sqrt{L^2 + \frac{\Delta A}{\tan\theta}} - L \tag{14}$$

In the case of a shrinking fold, the area of the smaller fold is:

$$\frac{a}{2} - \frac{\Delta A}{2} = \frac{1}{2}(L - \Delta L)^2 \tan\theta \tag{15}$$

We then find:

$$\Delta L = L - \sqrt{L^2 + \frac{\Delta A}{\tan\theta}} \tag{16}$$

#### Constant $h$ growth and shrinking

We first note:

$$L \tan\theta = h \tag{17}$$

$$(L + \Delta L) \tan(\theta - \Delta\theta) = h \tag{18}$$

We can write the equation for the area of the larger fold as:

$$\frac{a}{2} + \frac{\Delta A}{2} = \frac{1}{2}(L + \Delta L)^2 \tan(\theta - \Delta\theta) \tag{19}$$

Solving for $\Delta L$, we find that:

$$\Delta L = \sqrt{\frac{\frac{1}{2}L^2 tan(\theta) + \Delta A}{\tan(\theta - \Delta\theta)}} - L \tag{20}$$

In the case of a shrinking fold, the area of the smaller fold is:

$$\frac{a}{2} + \frac{\Delta A}{2} = \frac{1}{2}(L - \Delta L)^2 \tan(\theta + \Delta\theta) \tag{21}$$

Solving for $\Delta L$, we find that:

$$\Delta L = L - \sqrt{\frac{\frac{1}{2}L^2 tan(\theta) + \Delta A}{\tan(\theta + \Delta\theta)}} \tag{22}$$

While both constant $\theta$ and constant $h$ growth and shrinking could be at play in *T. adhaerens* perimeter folds, we observe striking cases of the former, where crease elongation is preceded by tissue action close to the outer edge of the fold (SI Movie 1).

### 5. Supplementary Figures

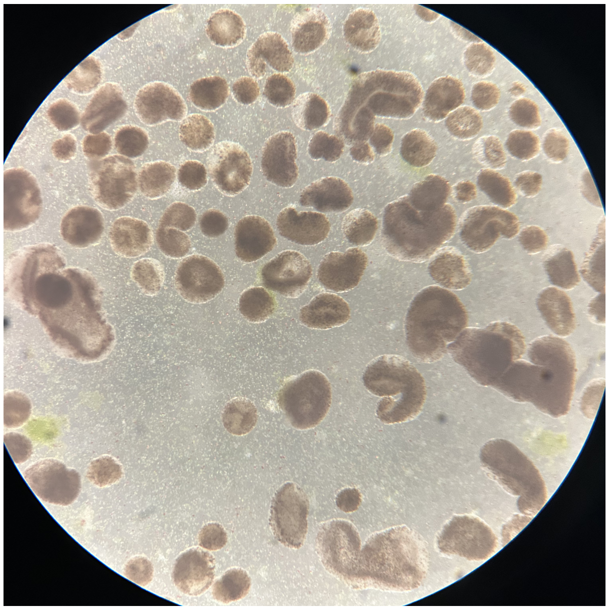

**SI Fig. S1**: **Detached, folded animals observed in culture dishes.** Image of laboratory culture of *T. adhaerens*. While most animals in culture are typically flat and adherent to the glass petri dish, we regularly observe a sub-population of detached animals which are typically in a folded state. This observation supports the intuition that substrate detachment is correlated with folding. What triggers folding in this case, when a rigid substrate is readily available, is not understood. Scale bar not available. Field of view is estimated to be $\sim 10mm \times 10mm$

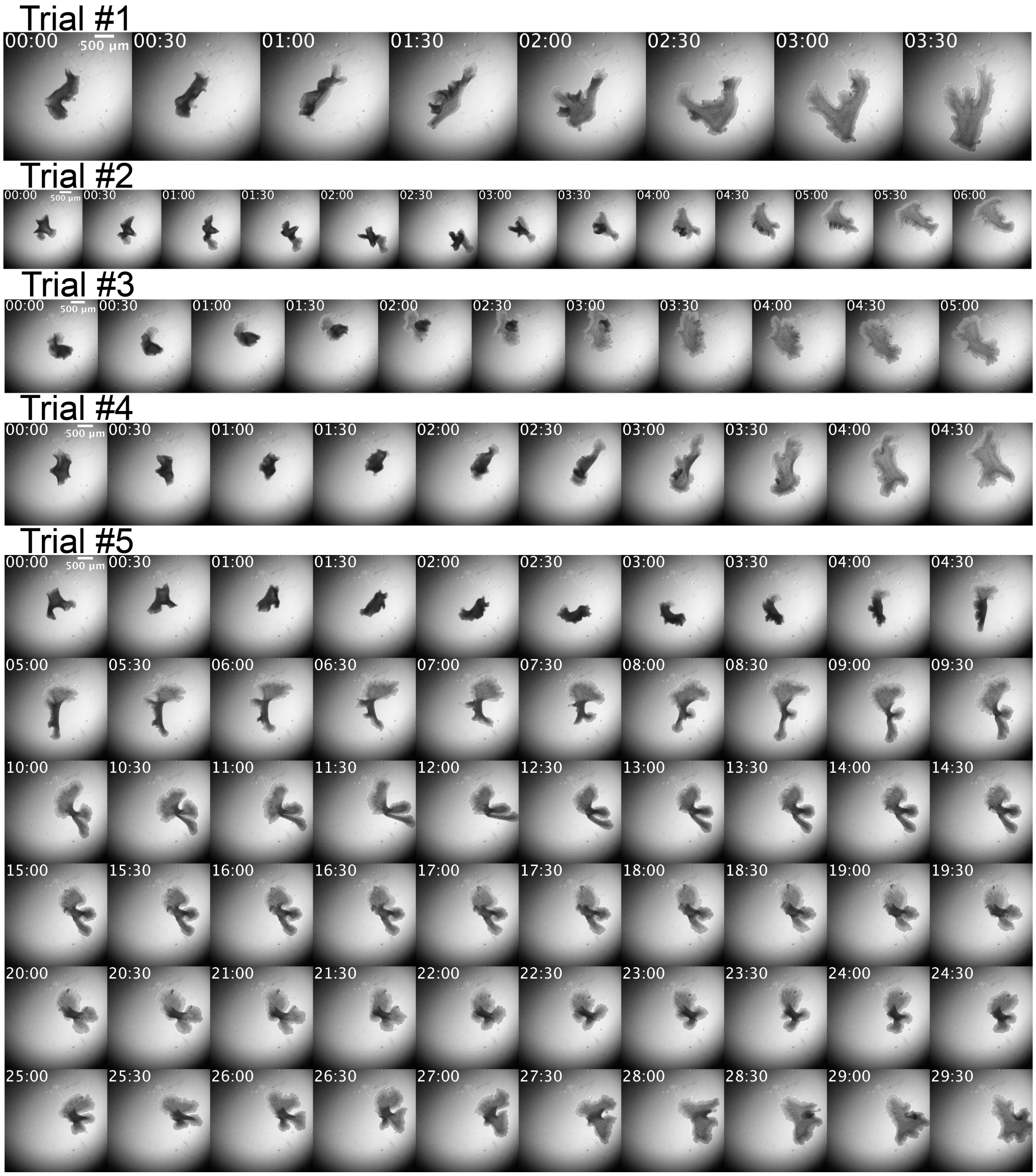


**SI Fig. S2**: **Repeated unfolding trials for the same animal.** A given animal repeatedly peeled from a rigid substrate performs characteristic unfolding behavior in an non-stereotypical fashion. In this case, the same animal took five very different paths to unfold in five repeated unfolding trials. Notably, in Trial 5 the animal passes through more complex folding states, which take longer to resolve. Scale bars are 500 $\mu m$.

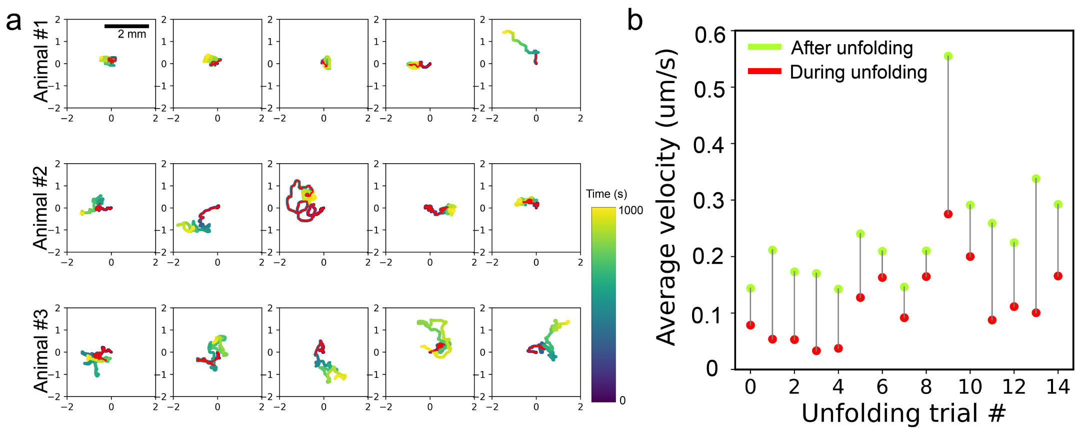


**SI Fig. S3**: **Animal trajectories during and after unfolding**. (a) Unfolding trajectories for five unfolding trials performed by three animals. Animal position was recorded during and after unfolding. Color map indicates time and red line indicates the portion of the track representing the unfolding trajectory. (b) Average animal velocity for fifteen unfolding trials shown in (a). In most cases, animal velocity is reduced during unfolding behavior.

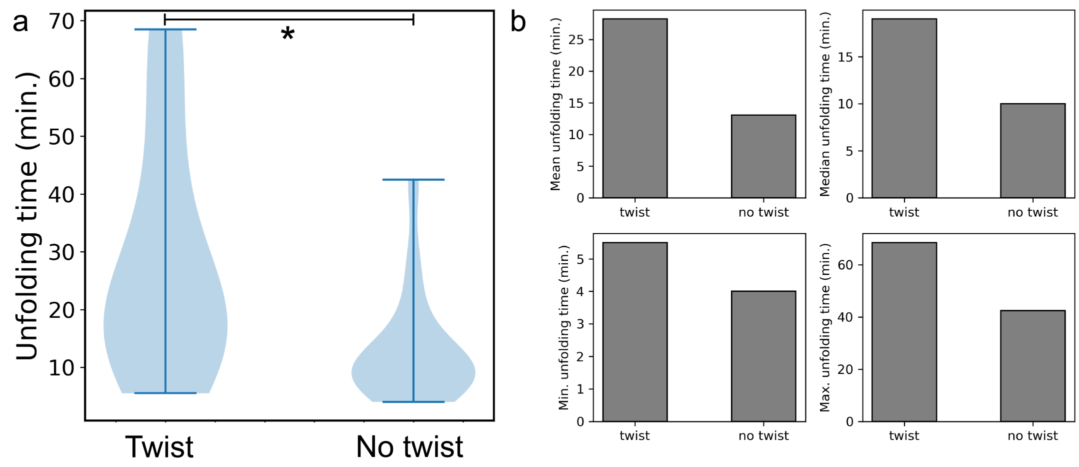


**SI Fig. S4**: **Unfolding time in large animals with and without $2\pi$ twist.** Ten repeated unfolding trials were performed for five large animals ($\sim$ 1 mm in diameter). These trials were then annotated based on whether or not a $2\pi$ twist was observed at any point during the unfolding trajectory. We observe that the presence of twist is associated with longer unfolding times (Welch's $t(16.7)=2.77$, p=0.013).

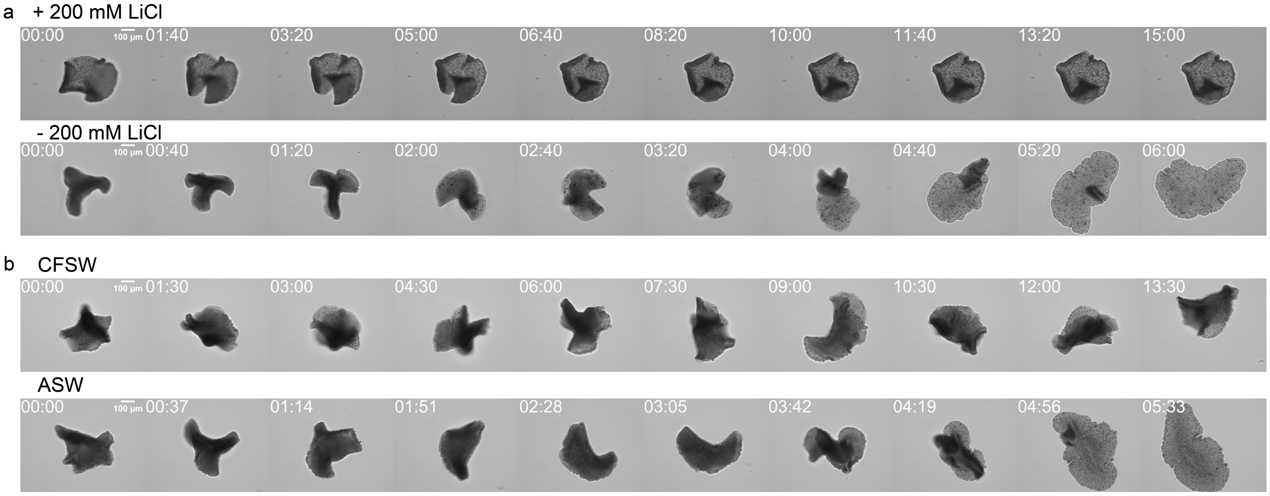


**SI Fig. S5**: **LiCl and CFSW reversibly inhibit unfolding behavior.** (a) Representative unfolding time lapse for a single animal unfolding with and without 200 mM lithium chloride (LiCl). (b) Representative unfolding time lapse of a single animal in artificial sea water and in calcium-free sea water (CFSW). In both cases, the treatment reversibly inhibits unfolding, which highlights the role of normal ciliary activity in driving unfolding. Scale bars are 100 $\mu m$.

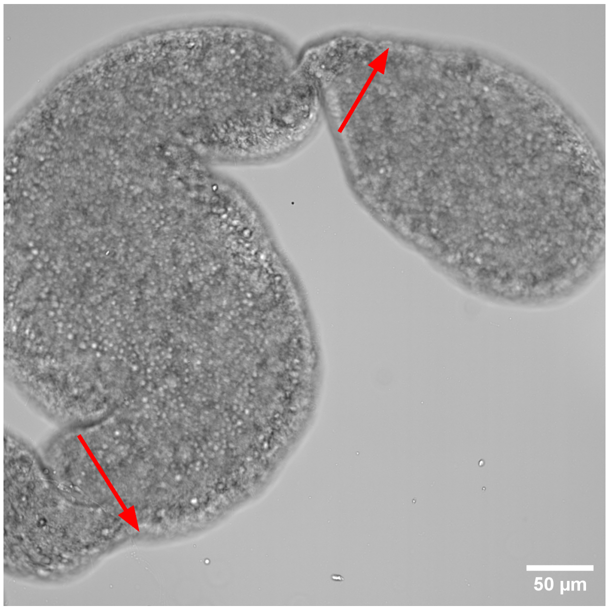


**SI Fig. S6**: **Two $2\pi$ twists with opposite chirality observed in the same animal.** Here a single animal exhibits two distinct $2\pi$ twists with opposite chirality, indicated with red arrows. In this configuration, twists can be removed through collision rather than being pushed out the edge of the animal. Scale bar is 50 $\mu m$.

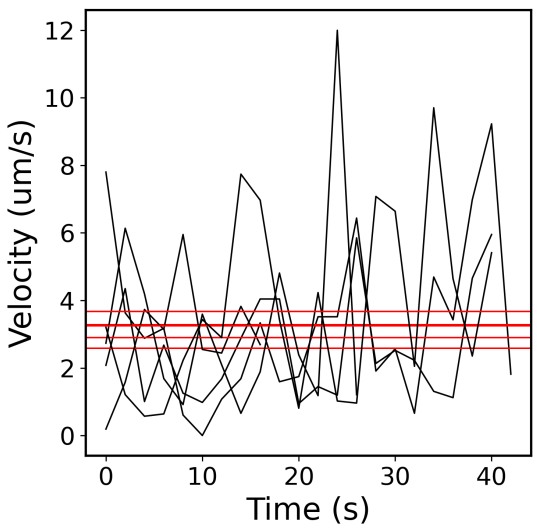


**SI Fig. S7**: **Average velocity of crease translation.** In crease datasets, crease translation events were identified and manually tracked. Translating creases move (in the animal frame of reference) at an average velocity of $\sim 3-4\ \mu m$.

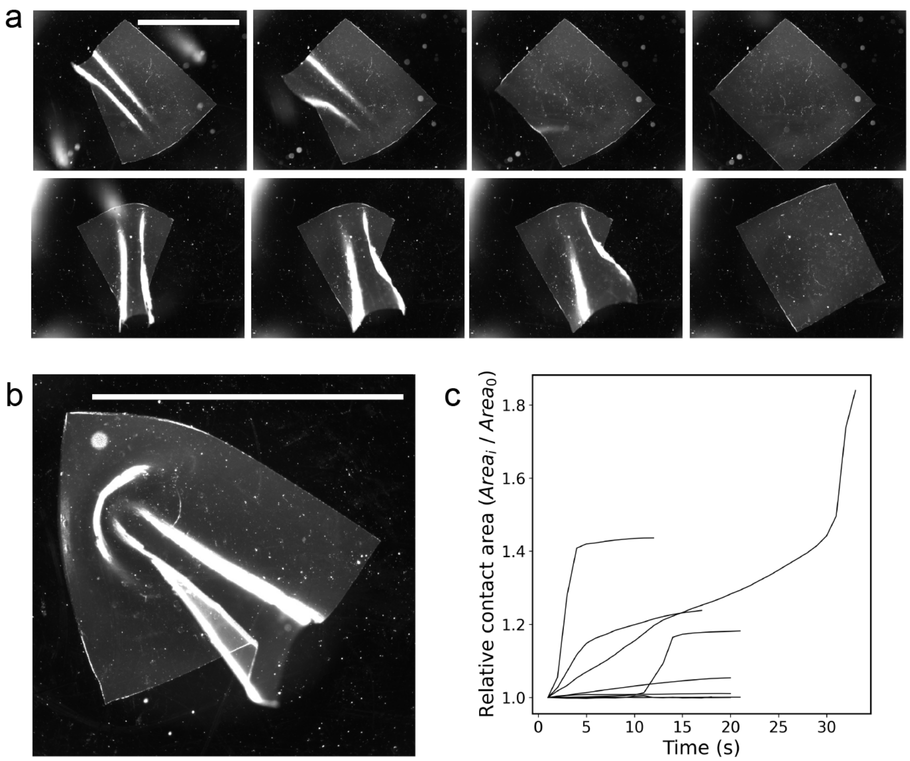


**SI Fig. S8**: **Unfolding behavior of thin rubber sheets.** (a) Unfolding behavior of thin silicone rubber sheets submerged in mineral oil on a glass surface. (b) Example of a frustrated folding state exhibited by a silicone rubber sheet. In this case, adhesion dominates over bending energy, leading to frustration. In the absence of activity, the sheet remains in a folded configuration. (c) Unfolding trajectories for several silicone rubber sheets. Notably, unlike in *T. adhaerens*, unfolding trajectories exhibit monotonically increasing contact area, and often show "snap"-like dynamics where bending energy drives unfolding.

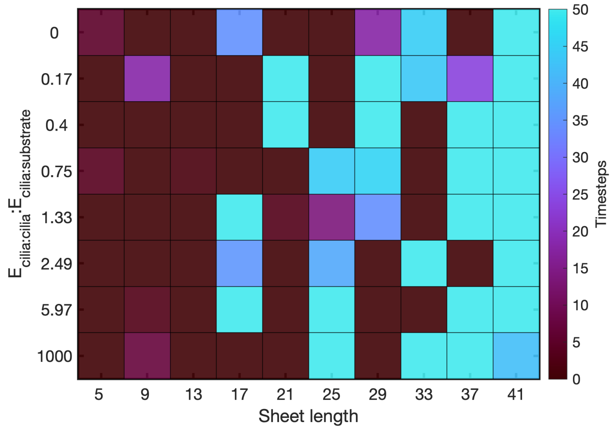


**SI Fig. S9**: **Impact of sheet size on unfolding time in 1D simulations.** In 1D simulations, we tested the impact of lattice length on unfolding time. We see that for the same single-fold initial condition, longer lattices exhibit longer unfolding times. This effect arises from the random probability that flat patches initiate new folds.

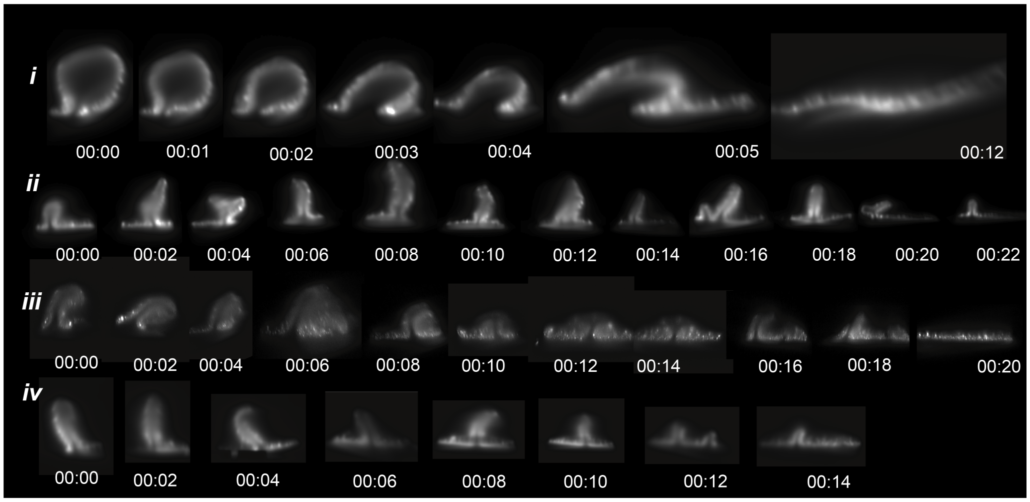


**SI Fig. S10**: **Ruck and hairpin fold cross sections.** Four fold cross sections extracted from 3D light sheet scans. Folds (i) and (ii) show higher temporal resolution of folds shown in Figure 2g. Folds (iii) and (iv) are additional hairpin folds used for analysis. In hairpin folds ii-iv, it is apparent that self-contact is exhibited until the fold is eventually removed.

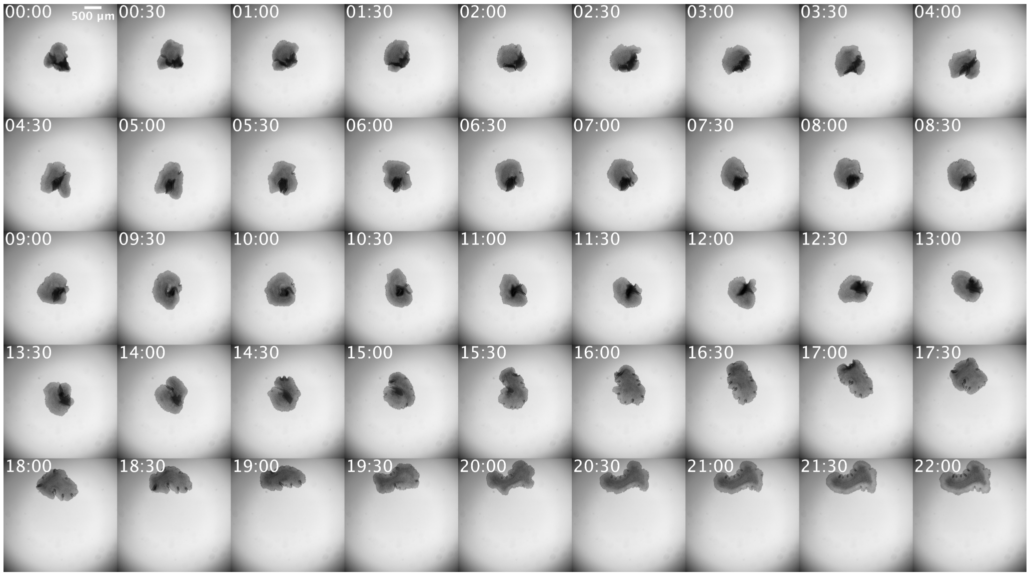


**SI Fig. S11**: **Upside down unfolding behavior.** An animal was detached from a substrate and allowed to settle in a dish. After a small amount of reattachment had occurred, the dish was sealed and inverted. We observe that normal unfolding still occurs against gravity. Scale bar is 500 $\mu m$.

### 6. Supplementary Movies

**Supplementary Movie 1**: Introduction to *T. adhaerens* unfolding behavior

**Supplementary Movie 2**: High-magnification imaging of live *T. adhaerens* folds

**Supplementary Movie 3**: Folded *T. adhaerens* observed in suspension via vertical tracking microscopy

**Supplementary Movie 4**: *T. adhaerens* folding on an ultra-thin glass capillary

**Supplementary Movie 5**: *T. adhaerens* folded state observed on colonial algae via light sheet microscopy

**Supplementary Movie 6**: Twisted *T. adhaerens* mid-unfolding body rupture

**Supplementary Movie 7**: Toroidal *T. adhaerens* unfolding trials

**Supplementary Movie 8**: Unfolding behavior in a ”stringy” animal

**Supplementary Movie 9**: 4D light sheet imaging of *T. adhaerens* unfolding behavior

**Supplementary Movie 10**: Fitting surfaces to 4D light sheet data

**Supplementary Movie 11**: Twist mobility and tightness modulation

**Supplementary Movie 12**: Ventral creases observed during unfolding behavior

**Supplementary Movie 13**: Ventral crease tip collisions

**Supplementary Movie 14**: Single cilia-resolved imaging of unfolding behavior

**Supplementary Movie 15**: Ciliary interaction across folds and twists

**Supplementary Movie 16**: Pinwheel pattern at the edge of a perimeter crease

**Supplementary Movie 17**: 2D simulation of unfolding via ciliary flocking
